# Timing and meaning in spoken thought relate to distinct patterns of brain dynamics

**DOI:** 10.64898/2026.11.18.767520

**Authors:** Donghui Song, Henyi Rao

## Abstract

When people think aloud, their reports vary in duration and in how closely successive thoughts are related in meaning. We asked whether these two dimensions relate to distinct patterns of brain dynamics. Using 10-minute think-aloud fMRI scans from 75 adults, we quantified timing as variability in sentence-event duration and semantic continuity as similarity between adjacent grouped thoughts. In models including both measures, greater duration variability was associated with lower relative brain entropy (BEN), estimated from BOLD time series, in the right dorsolateral prefrontal cortex (voxel wise FDR q < 0.05). This association persisted after adjustment for speech coverage, pause duration, articulation rate and event count. Semantic continuity was associated with differences in BOLD amplitude and synchrony across bilateral sensorimotor regions. These voxel wise associations fell below the FDR threshold after adjustment for speech coverage or pause duration. In a secondary atlas-based analysis, semantic continuity remained positively associated with local synchrony in Default C after adjustment for speech timing (β = 0.517, P = 0.003). These findings link the timing of spoken reports to dorsolateral prefrontal brain activity irregularity and show how the sensorimotor correlates of semantic continuity covary with speech timing.

## Introduction

A sentence can end while the thought it expresses continues. A speaker may develop one idea through several sentences of unequal length or shift to a new idea in a sentence much like the previous one in duration. Spoken thought thus has two measurable features: how much the duration of verbal reports varies and how closely successive thoughts are related in meaning. We asked whether individual differences in these features correspond to different properties of ongoing brain activity.

Dynamic accounts of spontaneous thought describe how internally generated experiences persist and change over time (Christoff et al., 2016). Neuroimaging studies have examined activity around reported thought onsets and transitions between inferred brain states (Ellamil et al., 2016; Tseng & Poppenk, 2020). Research on narrative comprehension has also shown that cortical activity reflects event structure at multiple timescales (Baldassano et al., 2017; Geerligs et al., 2022). Using think-aloud functional magnetic resonance imaging (fMRI), Su et al. (2025) related the boundaries and semantic organization of self-generated thoughts to activity and connectivity in default and control networks. Their recordings allow us to extend that work from the organization of individual events to differences between people across an entire sequence. When timing and meaning are measured in the same stream of spoken thought, which properties of the blood oxygen level-dependent (BOLD) signal are associated with each?

The answer may depend on which property of the BOLD signal is measured. Brain entropy (BEN), estimated here as the sample entropy of regional BOLD time series, characterizes temporal irregularity through the recurrence of similar patterns (Richman & Moorman, 2000; Wang et al., 2014). Fractional amplitude of low-frequency fluctuations (fALFF) quantifies the relative contribution of slow fluctuations. Regional homogeneity (ReHo) and voxel-mirrored homotopic connectivity (VMHC) describe synchrony among neighbouring voxels and between mirrored locations across hemispheres, respectively (Zang et al., 2004; Zou et al., 2008; Zuo et al., 2010). Associations between BEN and fALFF vary across regions at rest, and the two measures can respond differently within the same region under experimental modulation (Song et al., 2019; Song & Wang, 2026). Comparing these measures in the same recordings allows us to determine whether report duration variability and semantic continuity are associated with temporal irregularity, fluctuation amplitude or synchrony in different ways.

Thinking aloud also makes speech timing part of the scientific question. Intracranial recordings during speech production have identified distributed representations of semantic, articulatory and acoustic features (Verwoert et al., 2025). In spontaneous speech, elaborating an idea may coincide with longer stretches of speaking, while moving to another idea may coincide with a pause. An association between semantic continuity and BOLD activity may therefore share variance with the distribution of speech and silence. Speech coverage and pause duration characterize that distribution, while articulation rate describes how quickly words are produced during speech. The choice of temporal unit also matters. A single thought can span several sentences, so the variability of sentence durations and the variability of grouped-thought durations describe timing at different levels of the same sequence.

Here we analysed 10-minute think-aloud fMRI scans from 75 adults. We quantified temporal variability from sentence-event durations and semantic continuity from the similarity between adjacent grouped thoughts. We entered both measures in the same models to examine whether their associations differed in spatial distribution and across BEN, fALFF, ReHo and VMHC. We then examined how the semantic associations changed after adjustment for speech timing and whether the timing associations were sensitive to defining duration at the sentence or grouped-thought level. In secondary analyses, we examined seed-based functional connectivity (FC) and used joint BEN–FC models to estimate how much variation in report duration each measure accounted for after including the other.

## Results

### Behavioral timing and semantic continuity are related but distinct

The analyzed sample comprised 75 participants aged 18–36 years (mean 23.45, SD 4.27; 43 female and 32 male participants in the demographic record). Greater duration coefficient of variation (duration CV) accompanied lower adjacent semantic similarity (Pearson r = −0.310, two-sided P = 0.00676). The measures shared approximately 9.6% of their variance (Fig. 1c). Duration CV summarized sentence-level transcript events, whereas semantic similarity summarized adjacent grouped thoughts (Fig. 1a, b). Event durations and embedding similarities were independently recalculated for all 75 participants and matched the saved analysis values (Supplementary Table 1).

**Figure 1.**
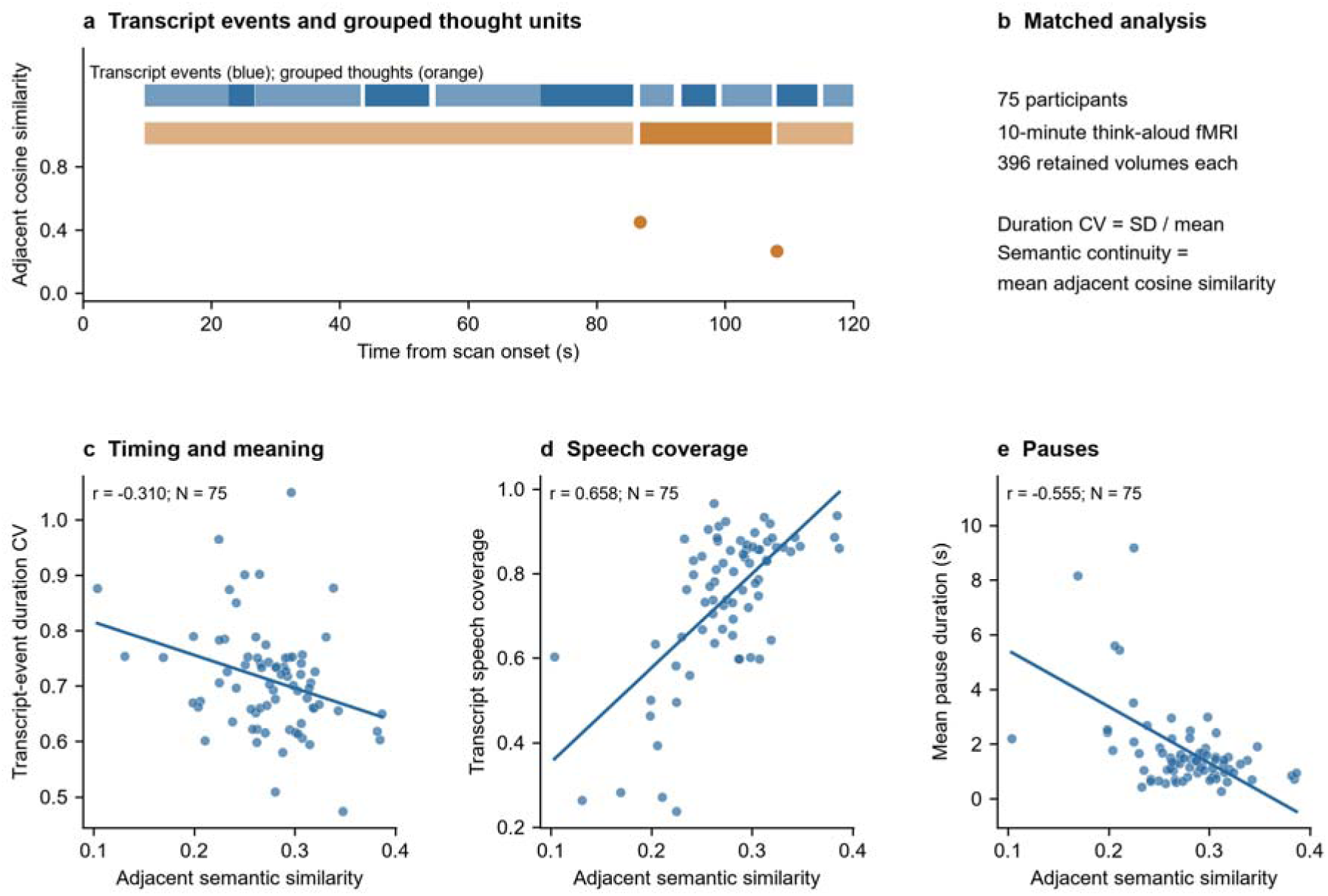
Temporal and semantic features of spoken thought. a, an observed example from sub-001 during the first 120 s. blue bars mark sentence-level transcript events; orange bars mark grouped thought intervals. Orange points show similarity to the previous grouped thought, positioned at the next thought’s onset. Duration CV uses the blue intervals, whereas semantic continuity uses adjacent grouped-thought embeddings. b, Matched analysis sample and measure definitions. c–e, Participant-level relationships between adjacent semantic similarity and transcript-event duration CV (c), speech coverage (d), and mean positive pause duration (e). Each point is one participant (N = 75); lines are ordinary least-squares fits. Pearson r values are shown; two-sided P values are 0.00676, 1.42 × 10□¹□ and 2.43 × 10□□, respectively. These descriptive correlations are unadjusted. Source data are provided.

Semantic continuity was also related to transcript-derived speech timing. Greater similarity between adjacent thoughts accompanied greater speech coverage (r = 0.658, P = 1.42 × 10□¹□) and shorter mean pauses (r = −0.555, P = 2.43 × 10□). We therefore assessed regional associations both before and after accounting for the measured distribution of speech and pauses (Fig. 1d, e).

### Sentence-level timing variability maps to lower relative prefrontal BEN

Greater duration CV was associated with lower BEN after adjustment for semantic continuity, age, sex and mean framewise displacement (FD). Twenty voxels survived Benjamini–Hochberg voxel wise false discovery rate (FDR) correction at q < 0.05 across 167,023 tested voxels. These formed two clusters of 13 and 7 voxels, with peaks at MNI coordinates (27.5, 24.2, 54.0), t (69) = −5.632, and (21.7, 41.6, 42.0), t (69) = −5.289. The corresponding uncorrected P-value cutoff selected by FDR was 5.36 × 10□. Atlas assignment placed 13 significant voxels in right Default A dorsal prefrontal parcels and 7 in a right Control B lateral dorsal prefrontal parcel (Fig. 2a, b; Supplementary Table 2). The selected-region scatter summarizes the direction of this association descriptively (Fig. 2c).

**Figure 2.**
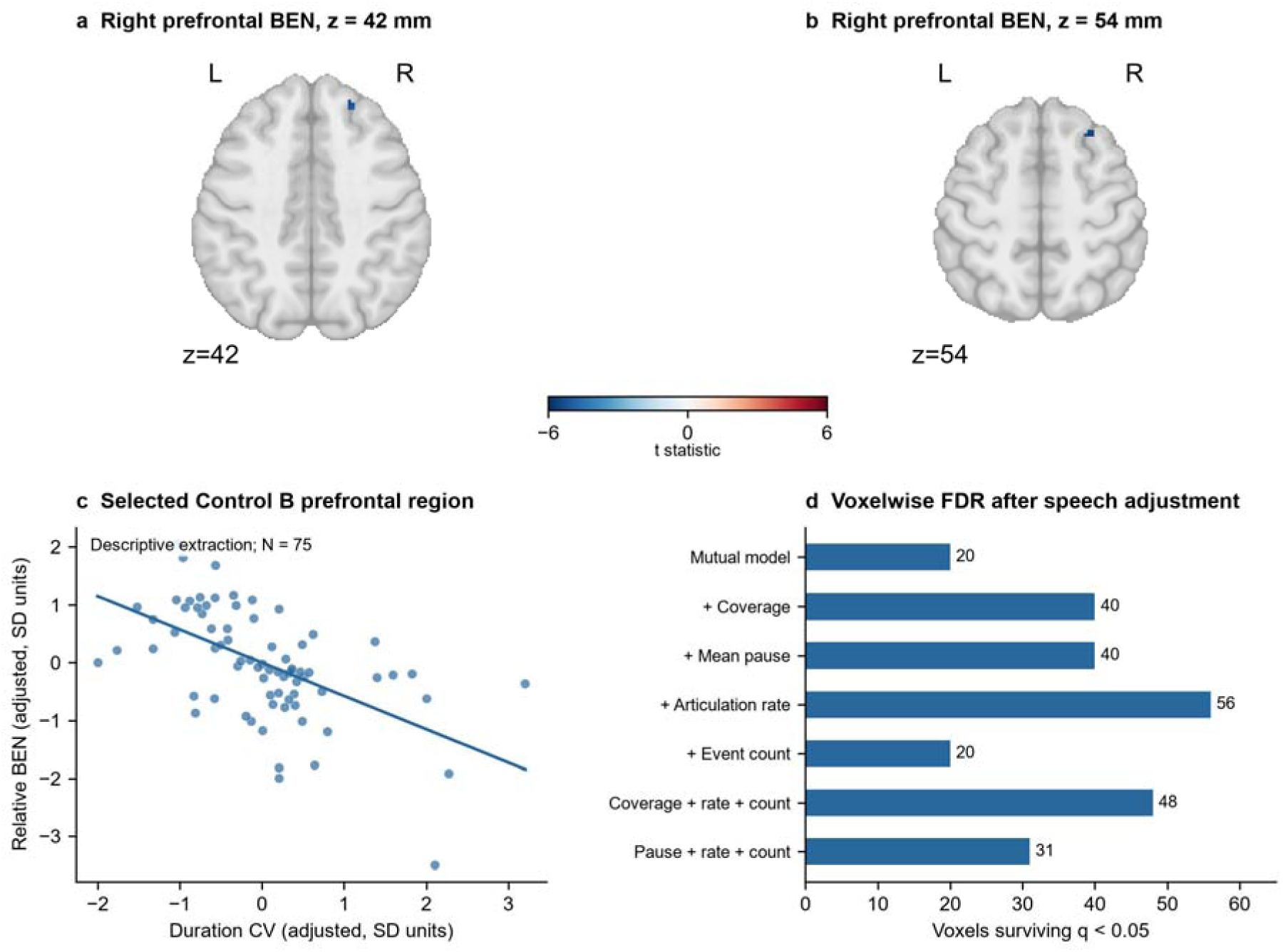
Sentence-event duration variability and relative prefrontal BOLD sample entropy. a, b, Axial views of the actual two-sided duration-CV coefficient map after voxel wise BH-FDR q < 0.05 across 167,023 voxels. Models include adjacent semantic similarity, age, sex and mean FD (N = 75; df = 69). Blue denotes a negative coefficient; the colour scale shows t values. Maps underwent grey-matter-confined, mask-normalized 8-mm smoothing and within-participant grey-matter centring. c, Descriptive partial-residual plot for the seven significant voxels assigned to Control B: both variables were standardized and residualized against the other behavioral measure, age, sex and FD. The fitted line is not an independent validation of the selected region. d, Numbers of FDR-significant voxels after separate or combined speech-timing adjustments. Coverage, pauses, articulation rate and event count are defined in Methods. Source data are provided.

To characterize its cortical distribution independently of the voxel summary, we tested the sentence-duration coefficient across all 400 Schaefer parcels, controlling the same covariates and applying Benjamini–Hochberg correction across the 400 parcel tests. Four right prefrontal parcels showed negative associations (standardized β = −0.432 to −0.455, q = 0.0310–0.0335), spanning Default A, Default B and Control B subdivisions (Supplementary Table 13). This parcel wise result supports a right prefrontal localization while showing that the association is not confined to a single network label.

The prefrontal association persisted after adjustment for speech timing. Adding coverage, mean pause duration, articulation rate or transcript-event count separately retained 40, 40, 56 and 20 FDR-significant voxels, respectively. A combined model containing coverage, articulation rate and transcript-event count retained 48 voxels; replacing coverage with mean pause duration retained 31. Thus, the association between duration variability and relative prefrontal sample entropy remained detectable across the tested speech-timing specifications (Fig. 2d; Supplementary Table 3).

### Semantic continuity covaries with a broad sensorimotor profile

In the mutually adjusted models, greater semantic continuity was associated predominantly with lower low-frequency amplitude and synchronization in bilateral sensorimotor territories. FDR correction identified 1,670 significant voxels for fractional amplitude of low-frequency fluctuations (fALFF), 1,092 for regional homogeneity (ReHo) and 937 for voxel-mirrored homotopic connectivity (VMHC; Fig. 3; Supplementary Table 2). The corresponding ALFF analysis showed a closely overlapping spatial pattern (Dice = 0.866; Supplementary Fig. 2).

**Figure 3.**
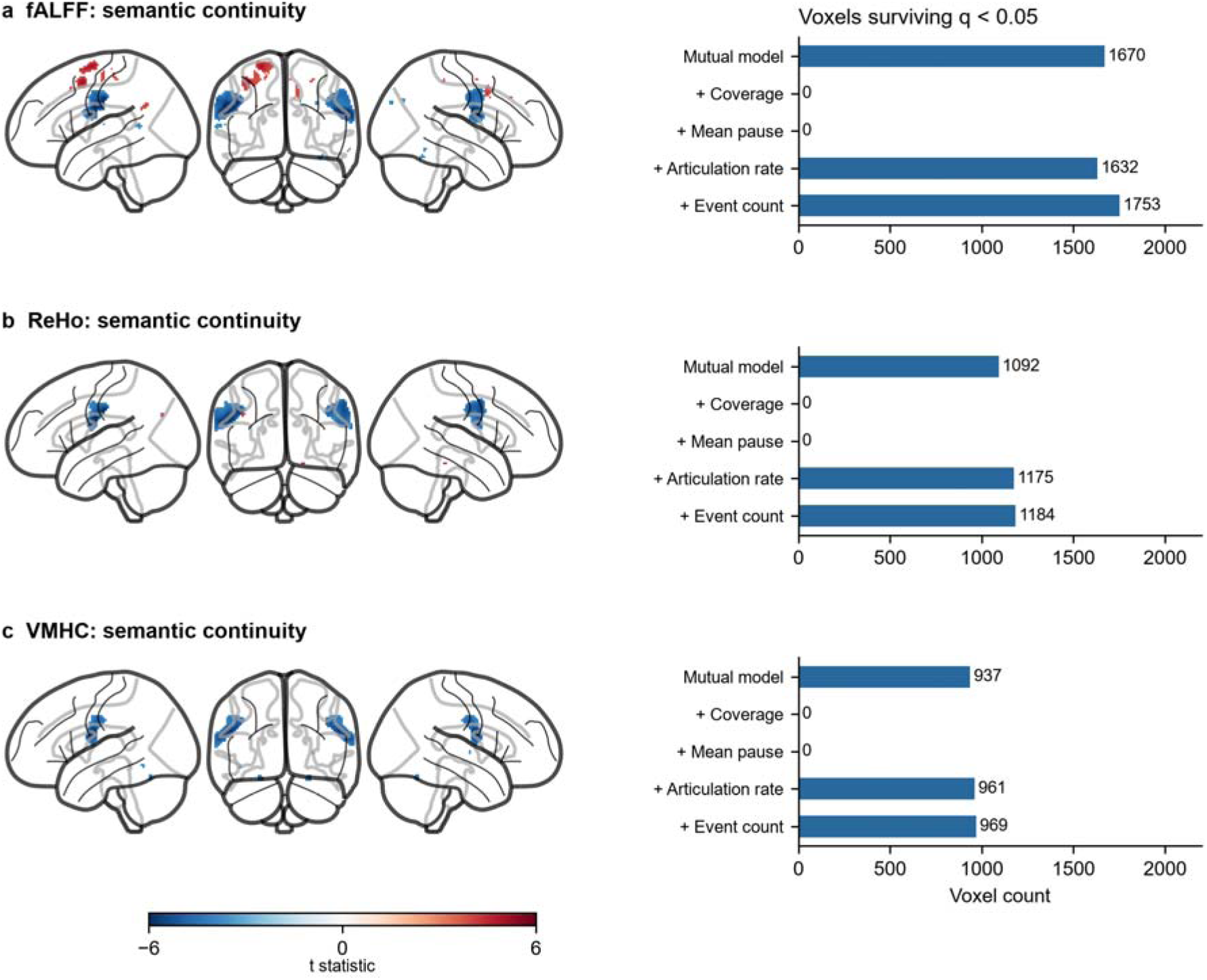
| Semantic continuity, regional BOLD amplitude and synchronization. a–c, Signed glass-brain projections of the actual FDR-thresholded semantic-similarity maps for fALFF, ReHo and VMHC. Blue and red indicate negative and positive coefficients, respectively; values outside the displayed ±6 t range are colour-saturated. All models include duration CV, age, sex and mean FD (N = 75; base df = 69). Maps use grey-matter-confined 8-mm smoothing and within-participant centring. The accompanying bars count significant voxels in the mutual model and after adding each speech-timing measure separately. Each model–metric map has its own voxel wise BH-FDR q < 0.05 correction; counts are descriptive summaries, not tests of differences between models. fALFF was calculated from unfiltered denoised series using the 0.009–0.1 Hz band; ReHo and VMHC used filtered series. Source data are provided. Corresponding ALFF maps and speech-timing adjustments are shown in Supplementary Fig. 2.

These associations changed substantially when speech coverage or pauses were included. Adding either coverage or mean pause duration left no voxels surviving FDR correction for fALFF, ReHo or VMHC. By comparison, adjustment for articulation rate or transcript-event count largely retained the associations: the corresponding numbers of significant voxels were 1,632 and 1,753 for fALFF, 1,175 and 1,184 for ReHo, and 961 and 969 for VMHC. The thresholded semantic maps were more sensitive to adjustment for speech coverage and pause duration than to adjustment for articulation rate or event count.

### Default C ReHo retains a conditional semantic association

Secondary regional analyses used atlas-defined Default A, Default C and Control B territories. These models included a speech-timing composite, articulation rate, transcript-event count, duration CV, age, sex and mean FD. Greater semantic continuity was associated with higher Default C ReHo (standardized β = 0.517, conventional 95% CI 0.180–0.854, nominal two-sided P = 0.00316, partial R² = 0.125). The association was supported by an HC3 heteroscedasticity-robust test (P = 0.00998) and 10,000 Freedman– Lane permutations (P = 0.0030).

Exploratory subdivision analysis identified positive semantic associations in the Default C inferior parietal region for ReHo (β = 0.499, conventional 95% CI 0.153–0.846, nominal P = 0.00536) and VMHC (β = 0.412, conventional 95% CI 0.082–0.741, nominal P = 0.0151). Their robust P values were 0.0216 and 0.0129, and permutation P values were 0.0059 and 0.0163, respectively. All three associations retained a positive coefficient when each participant was omitted in turn. These regional analyses used the complete valid atlas support and nominal P values; they are separate from the voxelwise FDR findings (Fig. 4; Supplementary Tables 4–6).

**Figure 4.**
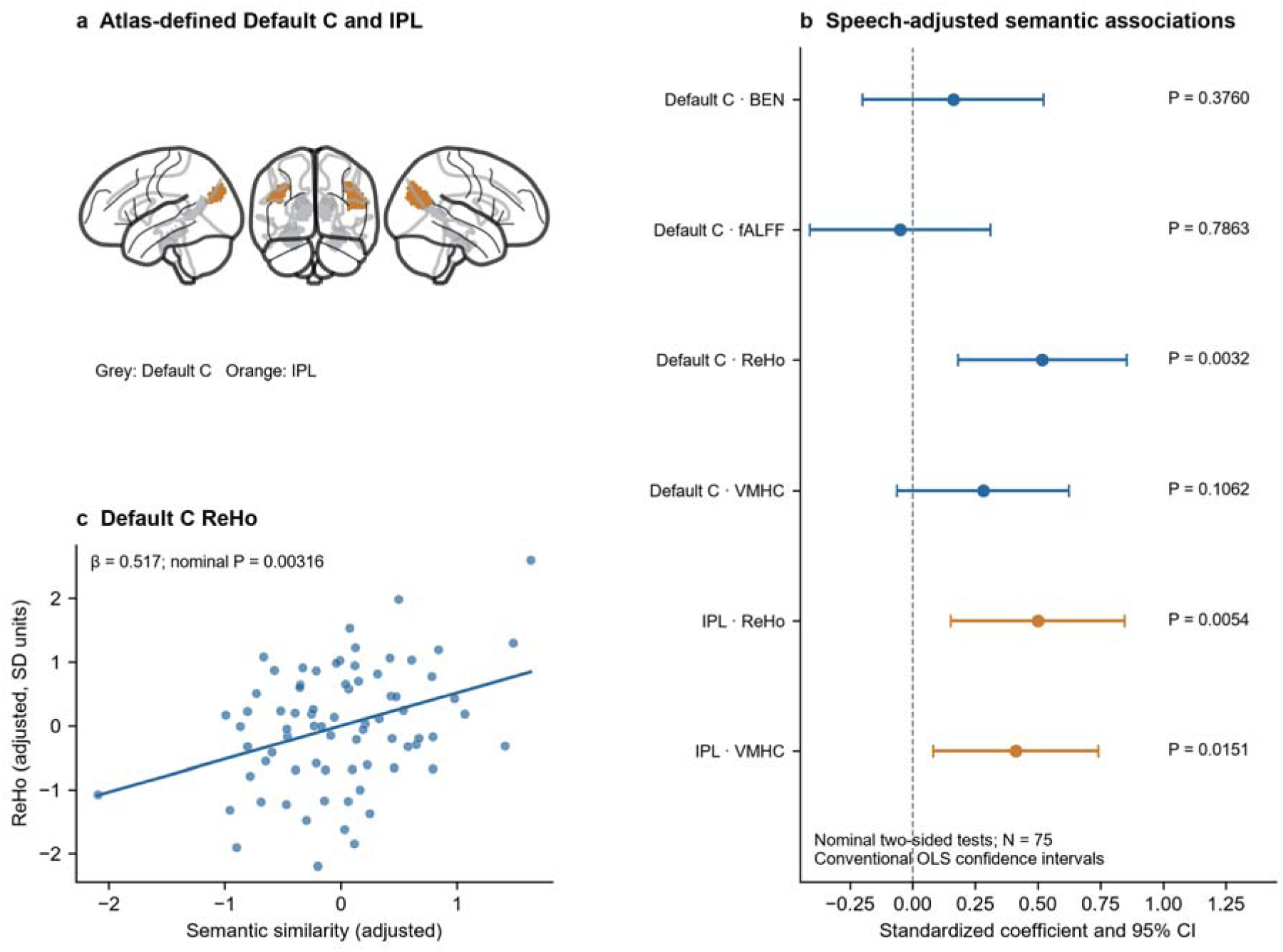
| Conditional semantic associations in cortical Default C. a, Full atlas-defined Default C support (grey) and its inferior parietal subdivision (IPL; orange), shown without an additional grey-matter restriction. b, Standardized semantic coefficients and conventional two-sided 95% t-reference confidence intervals for the four regional BOLD measures in Default C and the two reported exploratory IPL associations. N = 75; df = 66. All models include the speech-timing contrast, articulation rate, event count, duration CV, age, sex and mean FD. P values are nominal and are not corrected across ROIs or metrics. The complete cortical results appear in Supplementary Table 5. c, Partial-residual plot for Default C ReHo after removing the other predictors from standardized outcome and semantic similarity. Each point represents one participant; the line is an OLS fit. Robust and permutation tests are in Supplementary Table 6. Source data are provided.

### Timing associations across annotation levels

Replacing sentence-event duration CV with grouped-thought duration CV changed the temporal association. The two measures were weakly correlated (r = 0.205, 95% CI −0.023 to 0.413, P = 0.0779; Supplementary Fig. 1a). No BEN voxels survived FDR for grouped-thought duration CV in the mutual model or the speech-timing sensitivity models, including those that retained sentence-event count. In contrast, semantic associations remained FDR significant for fALFF (1,423 voxels), ReHo (923) and VMHC (563) in the mutual model. Adding coverage or mean pause again left no significant voxels (Supplementary Tables 9 and 10).

With sentence-event count held fixed, replacing duration CV retained the speech-adjusted semantic associations with Default C ReHo (β = 0.501, nominal P = 0.00243), IPL ReHo (β = 0.469, P = 0.00542) and IPL VMHC (β = 0.500, P = 0.00237). All three also remained detectable when grouped-thought count replaced sentence-event count (Supplementary Fig. 1b and Table 11). Exploratory transfer of the previous Control B seed to the grouped-thought outcome yielded no incremental BEN or connectivity association in the joint model (P = 0.551 and 0.916, respectively; Supplementary Table 12).

### Secondary analyses of seed connectivity and joint BEN–FC models

An approximate reanalysis of the original study’s connectivity approach recovered its reported association directions for thought-network clustering. Greater clustering accompanied stronger Control B–Default A connectivity (standardized β = 0.261, nominal P = 0.0235) and weaker within-Default C cortical connectivity (β = −0.252, nominal P = 0.0343), adjusting for age, sex and mean FD. Thought-network clustering summarizes the global semantic graph and is distinct from the adjacent semantic similarity used in the main models (Supplementary Table 7).

New seed analyses used the preprocessing matched to sample entropy. Greater duration CV was associated with weaker connectivity between the Control B prefrontal seed and the remaining voxels of Control B, conditional on semantic continuity, age, sex and mean FD (partial r ≈ −0.295; F (1,69) = 6.560, P = 0.0126; Fig. 5a, b). Direct correlations between BEN and seed-excluded connectivity were not detected: the unadjusted correlations were r = 0.145, P = 0.214 for Control B and r = 0.013, P = 0.911 for Default A.

**Figure 5.**
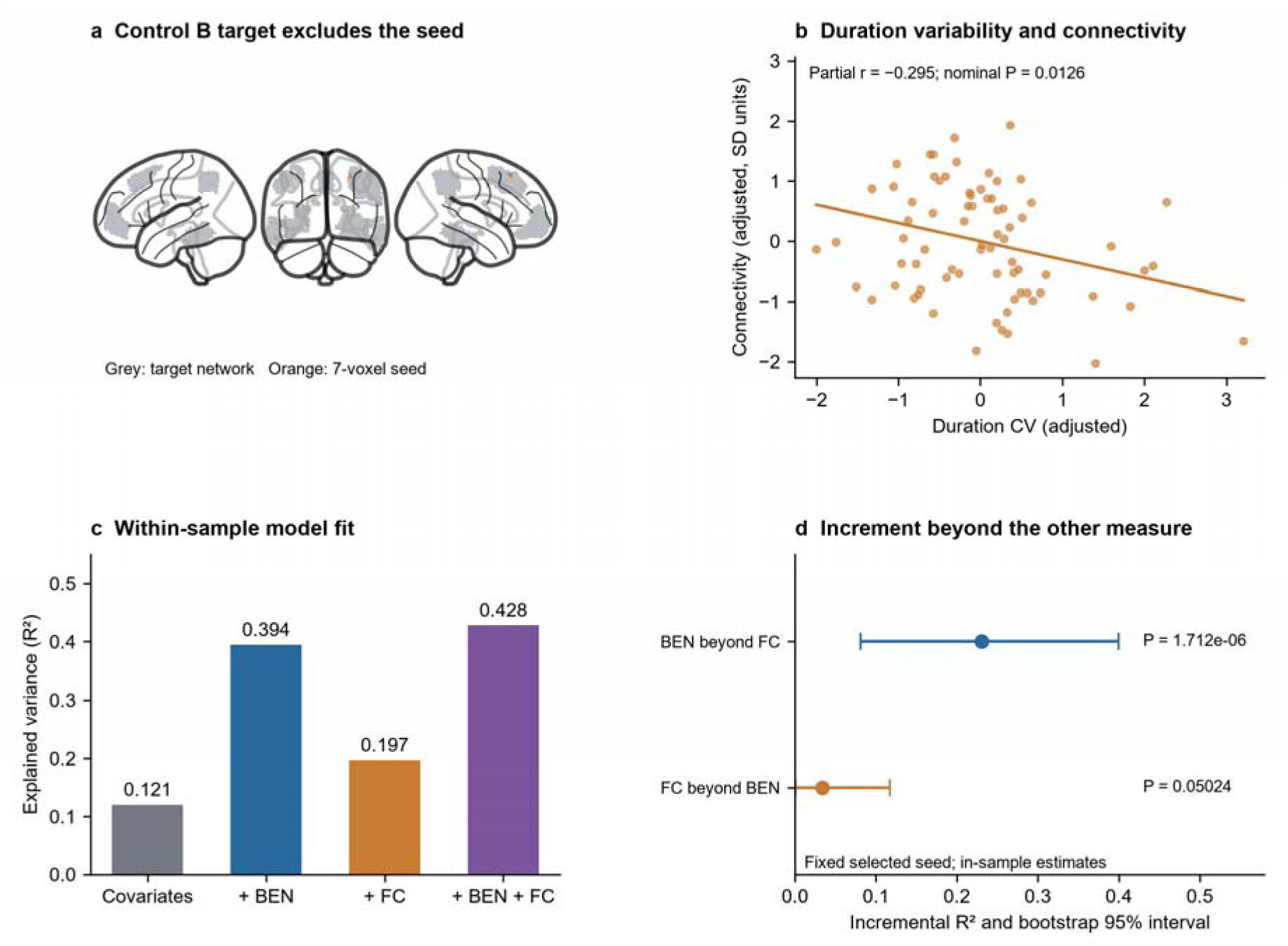
Secondary analysis of seed connectivity and joint BEN–FC models for sentence-event duration variability. a, the seven-voxel right prefrontal Control B seed (orange) and the remaining Control B target (grey). Connectivity correlates their voxel-mean series after excluding seed voxels from the target. b, standardized partial residuals for duration CV and Fisher-transformed connectivity, adjusting for semantic similarity, age, sex and FD (N = 75). The nominal P value comes from the corresponding nested F test (1,69 df). c, In-sample R² for the covariate model and additions of BEN, connectivity or both. d, In-sample incremental R² beyond the other imaging measure and covariates, with 95% percentile bootstrap intervals from 20,000 resamples. P values are nominal nested F tests (1,68 df). Bootstrap intervals for nonnegative ΔR² are not significance tests. Seeds were selected using duration CV in the same sample and held fixed during resampling. Full estimates appear in Supplementary Table 8. Source data are provided.

Nested regression models assessed the conditional associations of Control B prefrontal BEN and seed-excluded connectivity with duration CV. The base model (semantic continuity, age, sex and mean FD) had R² = 0.121; adding BEN yielded R² = 0.394, adding connectivity yielded R² = 0.197, and including both yielded R² = 0.428. Adding BEN beyond FC yielded ΔR² = 0.231 (nominal P = 1.71 × 10□). Adding FC beyond BEN yielded ΔR² = 0.033 (nominal P = 0.0502). These estimates describe in-sample fit for a seed selected using duration CV in the same cohort (Fig. 5c, d; Supplementary Table 8).

### Cross-validation defines the scope of the BEN prediction result

Repeated five-fold cross-validation clarified the scope of this result. With feature selection performed within each training fold and a BH-FDR threshold, BEN did not improve mean held-out Q² over the covariate model (mean ΔQ² = −0.001 across 20 repeats; range −0.092 to 0.163). A fixed training-fold p < 0.001 feature threshold, introduced after examining the FDR-based cross-validation, gave a mean ΔQ² of 0.080 (range −0.017 to 0.188; positive in 19/20 repeats). This exploratory gain was uneven across participants (40/75 had positive held-out error contributions; the largest contribution accounted for about 80% of the net gain) and voxel selection was spatially variable (median Jaccard = 0.258). Thus, the repeated split results do not establish a stable, generalizable prediction effect (Supplementary Tables 14 and 15).

As an exploratory comparator, BEN averaged across the four right lateral dorsal prefrontal Control B atlas parcels gave a mean held-out ΔQ² of 0.164 (range 0.110 to 0.210). This estimate describes repeated splits of the same cohort for an atlas region motivated by the study’s prefrontal findings (Source Data for Supplementary Table 14).

## Discussion

When people think aloud, the duration of their reports and the continuity of their meaning capture different aspects of the same sequence. In this study, greater variability in sentence-event duration was associated with lower relative BEN in right prefrontal cortex. Greater semantic continuity was associated with a broader pattern of BOLD amplitude and synchrony in bilateral sensorimotor cortex that was sensitive to speech coverage and pauses. A secondary atlas analysis also identified a more localized association between semantic continuity and regional homogeneity in Default C. Together, these findings extend work that has linked the boundaries and semantic organization of self-generated thought to default and control network activity (Su et al., 2025). They show how two features of the same stream of spoken thought relate to different properties of ongoing BOLD activity.

### Variation in report duration and prefrontal BEN

The association between sentence-event duration variability and right prefrontal BEN survived voxel wise correction and remained detectable after adjustment for semantic continuity and across models accounting for speech coverage, pauses, articulation rate and event count. A parcel wise analysis in the same cohort placed the association in neighboring prefrontal parcels spanning Default A, Default B and Control B. The convergence across spatial summaries supports a right prefrontal localization. Because BEN was centred on each participant’s grey-matter mean, the direction of the association is relative: participants with more variable report durations showed lower prefrontal BOLD sample entropy relative to their own grey-matter average. In this setting, lower BEN indicates greater regularity of recurring patterns in the local BOLD time series (Richman & Moorman, 2000; Wang et al., 2014). The association links variability in the duration of verbal reports across a scan to the temporal organization of regional BOLD fluctuations over the same scan.

The annotation scale helps specify the behavioral meaning of this association. Sentence-event duration variability was only weakly related to duration variability calculated for grouped thoughts, and the grouped-thought analysis yielded no BEN voxels surviving FDR correction. Grouped thoughts can span several sentences and include the pauses between them, so their duration summarizes a different feature of the sequence. Prior studies have localized changes in brain activity around reported thought onsets and inferred transitions (Ellamil et al., 2016; Tseng & Poppenk, 2020). The present result addresses a complementary level of description: how unevenly time is distributed across a person’s sentence-level reports. The sensitivity to annotation scale places the finding at the level of verbal expression and leaves open how moment-to-moment changes in prefrontal BOLD activity align with individual thought boundaries.

### Semantic continuity, speech timing and local synchrony

Across participants, greater similarity between adjacent thoughts accompanied greater speech coverage and shorter pauses. It was also associated predominantly with lower fALFF, regional homogeneity (ReHo) and interhemispheric homotopy in bilateral sensorimotor cortex. These voxelwise associations remained broadly detectable after adjustment for articulation rate or event count, whereas none survived FDR correction after adjustment for either speech coverage or mean pause duration. This contrast is informative because articulation rate describes how quickly words are produced during speech, while coverage and pause duration describe how speech and silence are distributed across the sequence. The pattern is consistent with shared variance between semantic continuity and the continuity of verbal output. Intracranial studies of speech production have identified distributed semantic representations alongside articulatory and acoustic representations (Verwoert et al., 2025). Our findings extend this question to spontaneously generated speech by showing that the sensorimotor BOLD profile associated with continuity of meaning depends on how the temporal organization of speaking is accounted for.

The Default C result adds a more localized observation. In an atlas-defined analysis, greater semantic continuity was associated with higher ReHo after adjustment for the transcript-derived speech measures and other covariates. The association also remained detectable when the timing covariate was recalculated at the grouped-thought level. ReHo indexes concordance among neighbouring BOLD time series, making this result relevant to local coordination within a functionally differentiated default network (Zang et al., 2004; Andrews-Hanna et al., 2010; Braga & Buckner, 2017). As a secondary ROI finding evaluated, it identifies a candidate regional correlate of semantic continuity conditional on measured speech timing. The broader sensorimotor and more localized Default C associations thus characterize different spatial aspects of how meaning is expressed during continuous speech.

### Local BEN and network connectivity

The seed analysis asked whether the timing of verbal reports also related to functional connectivity within the network containing a prefrontal BEN seed. Greater sentence-event duration variability was associated with weaker connectivity between the right prefrontal Control B seed and the rest of Control B. In the joint model, adding BEN to the connectivity model increased the proportion of variance explained within the sample by 0.231. Adding connectivity to the BEN model increased it by 0.033. These secondary findings quantify the conditional contribution of the two measures for a seed selected from the duration association in the same cohort. They also illustrate the distinction that motivated the use of multiple BOLD measures. BEN describes the temporal structure of a local BOLD series, while seed connectivity describes its covariation with a distributed signal. Prior work has shown that BEN and fALFF can relate differently across brain regions and respond differently within a region under experimental modulation (Song et al., 2019; Song & Wang, 2026). Cross-validation gains varied with the rule used to select voxels and were uneven across participants. The joint model therefore provides an estimate of within-sample fit for the selected region.

Speech timing was reconstructed from sentence-level transcript intervals because raw audio waveforms were unavailable. These intervals identify speech coverage and pauses between annotated events, while finer features such as brief within-sentence pauses, pitch and intensity require synchronized audio. Audio and event-aligned fMRI would allow future studies to examine how changes in meaning, pauses and vocal expression align around individual transitions.

Spoken thought unfolds through both the timing of its expression and the continuity of its meaning. In these recordings, variability in sentence-event duration was associated with relative right prefrontal BEN, while semantic continuity was associated with sensorimotor amplitude and synchrony that varied with the distribution of speech and pauses. A more focal association with Default C ReHo remained after speech timing was taken into account. Measuring timing and meaning together offers a way to relate the unfolding of spoken thought to both the temporal organization of local BOLD activity and its coordination across regions.

## Methods

### Participants, task and source data

We analyzed OpenNeuro ds006067 and the released annotations and embeddings accompanying the original study (Su et al., 2025). Applying the original imaging exclusion list to 118 available participants removed 39 for mean framewise displacement (FD) > 0.5 mm, two for anatomical anomalies, one for an imaging artifact and one for projector disconnection. The resulting 75 participants were used throughout. Participants spoke their thoughts aloud during a 10-minute fixation scan. Age and sex came from the participant file. The original procedures were approved by the Johns Hopkins Medicine and Homewood institutional review boards, with informed consent (Su et al., 2025).

### Temporal, semantic and speech-timing measures

Duration CV was the sample standard deviation (ddof = 1) divided by the mean of all duration entries in each participant’s onset-sorted BIDS event file. These entries matched sentence-level transcript intervals in all 75 participants. Event count was the number of these rows; the historical variable name “thought_count” in analysis files refers to this event count. Separately, sentence annotations were grouped by thoughtID to match the released embeddings. Semantic continuity was the mean cosine similarity between adjacent embeddings, originally generated with all-mpnet-base-v2 in Sentence Transformers 2.2.0 (768 dimensions) (Su et al., 2025). Thus, the temporal and semantic measures use different annotation levels. Supplementary Table 1 documents their correspondence and numerical verification.

Speech coverage was the total duration of merged, overlapping or touching sentence intervals divided by the span from first onset to last offset. Mean pause duration was the mean strictly positive gap between merged intervals (zero if none). Articulation rate was transcript word count divided by merged speech duration in minutes. Voxelwise sensitivity models added these measures or event count separately, or combined coverage or pause duration with rate and count. ROI models used T = z[(z(coverage) − z(mean pause))/√2], together with rate and count. This is an explicit standardized contrast, although some source files label it “speech temporal PC1”. Supplementary Methods provide interval handling and tokenization details.

### Acquisition and denoising

Acquisition used a 3-T Philips Ingenia Elition scanner: TR 1.5 s, TE 30 ms, flip angle 52°, slice thickness 2 mm, multiband factor 4 and j− phase encoding. Analyses began with distributed MNI152NLin2009cAsym derivatives from fMRIPrep 23.2.1 (Esteban et al., 2019). Removing four initial volumes left 396 per participant. Nuisance terms were six motion parameters, their derivatives, the squares of both sets (24 terms), and white-matter and cerebrospinal-fluid means. Constant and linear trends were removed and voxel series were scaled by their sample standard deviation. No global signal regression, volume censoring or spatial time-series smoothing was used. Motion control is relevant to both local and connectivity measures (Power et al., 2012; Ciric et al., 2017), and mean FD was included in group models.

Matched filtered and unfiltered series were generated. The former used a fifth-order forward–backward Butterworth filter at 0.009–0.1 Hz, with signal and confound filtering handled consistently in nilearn.signal.clean before regression. Matching their spectral treatment avoids the mismatch described in nuisance-regression studies (Hallquist et al., 2013). The latter retained the same nuisance terms and detrending without bandpass filtering. This filter was introduced in the present analysis.

### Regional BOLD measures

BENtbx estimated sample entropy from filtered series with embedding dimension m = 3 and tolerance r = 0.6 of within-voxel temporal standard deviation; no additional initial volumes were removed. Native maps were divided by 1,000 to recover sample-entropy units (Wang et al., 2014; Richman & Moorman, 2000). ALFF and fALFF used unfiltered denoised series: Fourier amplitude was the one-sided transform modulus divided by sample count; ALFF was its mean within 0.009–0.1 Hz and fALFF the sum in that band divided by the sum over all positive-frequency bins (Zou et al., 2008). ALFF therefore refers to amplitude in temporally standardized units.

ReHo was Kendall’s concordance among temporal ranks in a 3 × 3 × 3 neighbourhood of filtered series, calculated before smoothing (Zang et al., 2004). VMHC was the Fisher-transformed Pearson correlation between filtered series at a voxel and its nearest reflected MNI x-coordinate counterpart (Zuo et al., 2010). Both voxels had to be valid. No additional symmetric-template registration was applied. Exact edge, tie and numerical handling are given in Supplementary Methods.

### Spatial normalization of metric maps

Voxelwise maps used a resampled 2-mm MNI grey-matter probability template thresholded at 0.20. Mask-normalized 8-mm Gaussian smoothing divided the smoothed masked metric by its smoothed valid support, retaining support > 0.05. Each participant’s valid grey-matter mean was then subtracted, without division by spatial standard deviation. Group coverage required valid original-map values in at least 90% of participants. Atlas ROI maps used analogous smoothing and centring over complete valid cortical atlas support, without an external grey-matter threshold. ROI values were voxel-weighted means. These reference means were kept distinct.

### Voxelwise and regional inference

Voxelwise ordinary least squares included duration CV, semantic similarity, age, sex and mean FD plus an intercept (base residual df = 69). Continuous predictors were standardized using sample standard deviations; the female indicator was centred. Outcomes were spatially centred maps. Two-sided coefficient tests used Benjamini–Hochberg FDR q < 0.05 across voxels separately for each metric–behaviour map (Benjamini & Hochberg, 1995). No pooled correction across metrics or sensitivity models was applied. Cluster sizes describe connected suprathreshold voxels. The main BEN contrast concerned duration CV; complementary amplitude and synchronization contrasts concerned semantic continuity. This presentation hierarchy does not imply preregistration.

Atlas analyses followed Schaefer 400-parcel, 17-network labels (Schaefer et al., 2018; Yeo et al., 2011). Default A, cortical Default C and Control B were selected for correspondence with the original report; subdivision analyses were exploratory. Semantic models included T, articulation rate, event count, duration CV, age, sex and FD (df = 66). Continuous predictors and ROI outcomes were standardized. Regional P values were nominal, without across-metric or across-ROI correction. Conventional coefficient intervals and partial R² = t²/ (t² + df) was supplemented by HC3 inference, 10,000 Freedman–Lane permutations and leave-one-participant-out coefficients for the reported associations. Definitions and full cortical results appear in Supplementary Tables 4–6. Separately, the sentence-duration CV coefficient was tested across all 400 available cortical parcels using parcel-mean BEN and the same covariates; BH-FDR q < 0.05 controlled the parcelwise spatial family. This was a complementary same-cohort localization analysis, not an independent replication.

### Connectivity and joint models

A contextual approximation of the original connectivity analysis used full-series regression of six motion parameters, global, white-matter and cerebrospinal-fluid signals and linear/quadratic trends, followed by exclusion of the first ten volumes and FD ≥ 1 mm frames with two neighbours on each side. No extra bandpass filter was used. Correlations between bilateral atlas subregions were Fisher transformed and averaged; thought-network clustering models included age, sex and FD. This cortical implementation and its differences from the new analysis are detailed in Supplementary Methods and Table 7.

New connectivity used the BEN-matched filtered series. Intersecting the 20 significant BEN voxels with network labels produced 13-voxel Default A and 7-voxel Control B prefrontal seeds. Each seed’s mean series was correlated with the voxel mean of its network after excluding the seed, then Fisher transformed. Joint models predicted standardized duration CV from semantic similarity, age, sex and FD, adding standardized Control B BEN, connectivity or both. Incremental R² used nested F tests, 20,000 participant bootstrap resamples and 10,000 reduced-model residual permutations. Seeds remained fixed during resampling: the resulting estimates concern within-sample conditional associations. Supplementary Methods and Table 8 give full algorithms, seeds and estimates.

### Repeated cross-validation

We used 20 repeated shuffled five-fold partitions of the 75 participants (60 training and 15 test participants per fold). The baseline included semantic similarity, age, sex and mean FD. Voxel selection used the training-fold two-sided duration-CV coefficient map, intersected with Control B after whole-grey-matter BH-FDR or uncorrected thresholding. Fixed thresholds (P < 0.001, 0.005 and 0.01) were examined after the FDR-based analysis. When selection was empty, the augmented model used covariate-only predictions for that fold. Feature scaling and model fits were estimated from training data. An exploratory fixed-atlas comparator used BEN averaged in the valid-grey-matter intersection of the four right lateral dorsal prefrontal Control B parcels. Held-out Q² was scored relative to fold-specific training-mean predictions. ΔQ² denotes the increment over the covariate model. Ranges summarize variation across splits.

### Temporal-unit sensitivity analysis

For each participant, we grouped sentence annotations by the released thoughtID and defined grouped-thought duration as the last sentence offset minus the first sentence onset, including intervening pauses. CV was the sample standard deviation divided by mean duration. Embedding counts and chronological order were checked against these groups. This CV replaced sentence-event CV in the same 75-participant models. To separate CV definition from count definition, count-adjusted models first retained sentence-event count; a further specification replaced it with the number of grouped thoughts. ROI contrasts followed the same sequence. Spatial processing and voxelwise correction were unchanged. With no FDR-significant grouped-thought BEN seed, the previous sentence-selected seeds were retained only for exploratory transfer analyses; no new seeds were defined from uncorrected maps. These analyses characterize sensitivity to operationalization, rather than a formal test of differences between temporal-level coefficients.

## Data availability

Source imaging and behavioural data are available from OpenNeuro, accession ds006067 (https://openneuro.org/datasets/ds006067). Released thought annotations, embeddings and original analysis materials are available at https://doi.org/10.5281/zenodo.15665444.

## Declaration of competing interest

The authors declare no competing interests.

## Notes

https://openneuro.org/datasets/ds006067

### Competing Interest Statement

The authors have declared no competing interest.

## References

Andrews-Hanna, J. R. et al. Functional-anatomic fractionation of the brain’s default network. Neuron 65, 550–562 (2010). 10.1016/j.neuron.2010.02.005

Baldassano, C. et al. Discovering event structure in continuous narrative perception and memory. Neuron 95, 709–721.e5 (2017). 10.1016/j.neuron.2017.06.041

Benjamini, Y. & Hochberg, Y. Controlling the false discovery rate: a practical and powerful approach to multiple testing. J. R. Stat. Soc. B 57, 289–300 (1995). 10.1111/j.2517-6161.1995.tb02031.x

Braga, R. M. & Buckner, R. L. Parallel interdigitated distributed networks within the individual estimated by intrinsic functional connectivity. Neuron 95, 457–471.e5 (2017). 10.1016/j.neuron.2017.06.038

Christoff, K., Irving, Z. C., Fox, K. C. R., Spreng, R. N. & Andrews-Hanna, J. R. Mind-wandering as spontaneous thought: a dynamic framework. Nat. Rev. Neurosci. 17, 718–731 (2016). 10.1038/nrn.2016.113

Ciric, R. et al. Benchmarking of participant-level confound regression strategies for the control of motion artifact in studies of functional connectivity. NeuroImage 154, 174–187 (2017). 10.1016/j.neuroimage.2017.03.020

Ellamil, M. et al. Dynamics of neural recruitment surrounding the spontaneous arising of thoughts in experienced mindfulness practitioners. NeuroImage 136, 186–196 (2016). 10.1016/j.neuroimage.2016.04.034

Esteban, O. et al. fMRIPrep: a robust preprocessing pipeline for functional MRI. Nat. Methods 16, 111– 116 (2019). 10.1038/s41592-018-0235-4

Geerligs, L. et al. A partially nested cortical hierarchy of neural states underlies event segmentation in the human brain. eLife 11, e77430 (2022). 10.7554/eLife.77430

Hallquist, M. N., Hwang, K. & Luna, B. The nuisance of nuisance regression: spectral misspecification in a common approach to resting-state fMRI preprocessing reintroduces noise and obscures functional connectivity. NeuroImage 82, 208–225 (2013). 10.1016/j.neuroimage.2013.05.116

Power, J. D. et al. Spurious but systematic correlations in functional connectivity MRI networks arise from subject motion. NeuroImage 59, 2142–2154 (2012). 10.1016/j.neuroimage.2011.10.018

Richman, J. S. & Moorman, J. R. Physiological time-series analysis using approximate entropy and sample entropy. Am. J. Physiol. Heart Circ. Physiol. 278, H2039–H2049 (2000). 10.1152/ajpheart.2000.278.6.H2039

Schaefer, A. et al. Local-global parcellation of the human cerebral cortex from intrinsic functional connectivity MRI. Cereb. Cortex 28, 3095–3114 (2018). 10.1093/cercor/bhx179

Song, D. & Wang, Z. Age-dependent effects of intranasal oxytocin administration were revealed by resting brain entropy (BEN). Behav. Brain Res. 500, 115985 (2026). 10.1016/j.bbr.2025.115985

Song, D., Chang, D., Zhang, J., Ge, Q., Zang, Y.-F. & Wang, Z. Associations of brain entropy (BEN) to cerebral blood flow and fractional amplitude of low-frequency fluctuations in the resting brain. Brain Imaging Behav. 13, 1486–1495 (2019). 10.1007/s11682-018-9963-4

Su, H., Li, X., Born, S., Honey, C. J., Chen, J. & Lee, H. Neural dynamics of spontaneous memory recall and future thinking in the continuous flow of thoughts. Nat. Commun. 16, 6433 (2025). 10.1038/s41467-025-61807-w

Tseng, J. & Poppenk, J. Brain meta-state transitions demarcate thoughts across task contexts exposing the mental noise of trait neuroticism. Nat. Commun. 11, 3480 (2020). 10.1038/s41467-020-17255-9

Verwoert, M. et al. Whole-brain dynamics of articulatory, acoustic and semantic speech representations. Commun. Biol. 8, 432 (2025). 10.1038/s42003-025-07862-x

Wang, Z., Li, Y., Childress, A. R. & Detre, J. A. Brain entropy mapping using fMRI. PLoS ONE 9, e89948 (2014). 10.1371/journal.pone.0089948

Yeo, B. T. T. et al. The organization of the human cerebral cortex estimated by intrinsic functional connectivity. J. Neurophysiol. 106, 1125–1165 (2011). 10.1152/jn.00338.2011

Zang, Y., Jiang, T., Lu, Y., He, Y. & Tian, L. Regional homogeneity approach to fMRI data analysis. NeuroImage 22, 394–400 (2004). 10.1016/j.neuroimage.2003.12.030

Zou, Q. H. et al. An improved approach to detection of amplitude of low-frequency fluctuation (ALFF) for resting-state fMRI: fractional ALFF. J. Neurosci. Methods 172, 137–141 (2008). 10.1016/j.jneumeth.2008.04.012

Zuo, X. N. et al. Growing together and growing apart: regional and sex differences in the lifespan developmental trajectories of functional homotopy. J. Neurosci. 30, 15034–15043 (2010). 10.1523/JNEUROSCI.2612-10.2010

