## Supplementary Materials for "Timing and meaning in spoken thought relate to distinct patterns of brain dynamics"

### Supplementary Methods

#### Participants and source data

We analysed OpenNeuro ds006067 and the released transcript annotations and embeddings accompanying the original study (Su et al., 2025). The local dataset documentation identifies the addition of fMRIPrep and FreeSurfer derivatives in release v2.0.0. The original imaging exclusion list was applied to the 118 publicly available participants: 39 were excluded for mean FD > 0.5 mm, two for anatomical anomalies, one for an imaging artifact and one for projector disconnection. This yielded the matched sample of 75 used throughout the reported analyses. Age and sex were taken from the participant file. The original procedures were approved by the Johns Hopkins Medicine and Homewood institutional review boards, with informed consent (Su et al., 2025). [AUTHOR TO COMPLETE: applicable institutional determination for the present public-data secondary analysis and the exact archived dataset snapshot identifier.]

#### Task and behavioural measures

Participants spoke their thoughts aloud during a 10-minute scan while viewing a fixation cross on a grey screen. Speech was recorded and transcribed in the original study. We used the distributed BIDS transcript events and sentence-level annotations without generating new segmentations. BIDS rows match the sentence annotations; grouped thoughtID units match the embedding rows. These are different temporal units. Duration CV was calculated from all event-duration entries in each participant’s BIDS events file after sorting by onset, using the sample standard deviation (ddof = 1) divided by the arithmetic mean. Transcript-event count was the number of event rows.

Semantic continuity used the released thought embeddings, generated with all-mpnet-base-v2 in Sentence Transformers 2.2.0 (768 dimensions) (Su et al., 2025). Embedding row counts were checked against the grouped thought annotations. For each participant, a cosine-similarity matrix was calculated and its first upper diagonal averaged, yielding the mean similarity of adjacent thoughts. No additional category restriction was applied in this calculation. This measure was distinct from the earlier TF–IDF exploratory measure and from average thought-network clustering used in the contextual connectivity analysis.

#### Transcript-derived speech timing

Sentence onset–offset intervals were sorted, and overlapping or touching intervals were merged. Invalid intervals and intervals with non-positive duration were excluded. Speech coverage was the summed merged duration divided by the span between the first onset and last offset. Mean pause duration was the mean strictly positive inter-interval gap, assigned zero if no positive gaps occurred. Articulation rate was transcript word count divided by merged duration in minutes. Words were counted using the regular expression \b[\w’']+\b. These quantities estimate speech timing from annotations, rather than detecting phonation from audio.

Voxelwise sensitivity models added coverage, mean pause duration, articulation rate or transcript-event count individually. Two combined models added coverage, articulation rate and transcript-event count, or mean pause duration, articulation rate and transcript-event count. ROI models used a composite T = z[(z(coverage) − z(mean pause))/√2], with sample-standard-deviation standardization across participants, together with articulation rate and transcript-event count. Although labelled “speech temporal PC1” in some intermediate files, the implemented quantity was this explicit standardized contrast.

#### Acquisition and initial preprocessing

Data were acquired on a 3-T Philips Ingenia Elition scanner at the Kennedy Krieger Institute. The distributed acquisition metadata specify TR = 1.5 s, TE = 30 ms, flip angle = 52°, slice thickness = 2 mm, multiband acceleration factor = 4 and j− phase encoding. Analyses began with the distributed MNI152NLin2009cAsym BOLD derivatives generated by fMRIPrep (version 23.2.1) (Esteban et al., 2019). These derivatives, rather than a newly executed fMRIPrep workflow, were the inputs to the present denoising. The acquisition and preprocessing descriptions refer to distributed derivatives; a new fMRIPrep run was not performed.

#### Matched denoising

Four initial volumes were removed before denoising, leaving 396 volumes for each of the 75 participants. Nuisance regression included the six rigid-body motion parameters, their first derivatives, the squares of the original parameters and derivatives (24 motion terms), and mean white-matter and cerebrospinal-fluid signals. Constant and linear trends were removed, and voxel series were standardized using their sample standard deviation. No global signal regression or spatial smoothing was applied in this pipeline.

Filtered and unfiltered outputs were generated. The filtered series used a fifth-order Butterworth filter applied forward and backward between 0.009 and 0.1 Hz. Signal and confound filtering was handled consistently within nilearn.signal.clean before nuisance regression. The unfiltered output retained the same detrending and nuisance terms without temporal bandpass filtering. Volumes were not censored, preserving contiguous samples for entropy estimation. Mean FD was included in group models. This 0.009–0.1 Hz filter was introduced in the present analysis, not inherited from the original connectivity analysis.

#### BOLD sample entropy

BEN was calculated with BENtbx from the filtered series using embedding dimension m = 3 and tolerance r = 0.6 of the within-voxel temporal standard deviation. No further dummy volumes were removed within the toolbox. Native maps, stored at 1,000 units per sample-entropy unit, were divided by 1,000 before downstream analyses. The pipeline included independent Chebyshev-distance pair-count checks at sampled voxels; these were numerical implementation checks rather than additional behavioural tests. We use BEN to denote BOLD sample entropy under the specified parameters (Wang et al., 2014; Richman & Moorman, 2000).

#### ALFF and fALFF

Both amplitude measures used the unfiltered denoised series to preserve the full frequency spectrum. After temporal mean removal, the one-sided real fast Fourier transform was computed. Amplitude at each frequency was the transform modulus divided by the number of samples. ALFF was the mean amplitude of bins from 0.009 to 0.1 Hz, inclusive. fALFF was the sum of amplitudes in those bins divided by the sum across all strictly positive-frequency bins, including the Nyquist bin when present; the denominator was bounded below at 10⁻¹². Because the input series had already been temporally standardized, ALFF describes band-limited amplitude in those standardized units, not raw scanner-signal amplitude (Zou et al., 2008).

#### ReHo and VMHC

ReHo was calculated before spatial smoothing using temporal ranks and Kendall’s coefficient of concordance within a 3 × 3 × 3 neighbourhood (Zang et al., 2004). Average ranks were assigned to ties. In-mask neighbours contributed at edges, with at least two required. For K available neighbours and N samples, W = 12 Σt(Rt − K(N + 1)/2)²/[K²(N³ − N)], where Rt is the summed rank at time t. The implemented denominator did not include a separate tie correction.

VMHC used the filtered denoised series. Voxel coordinates were transformed to MNI world coordinates, the x coordinate was reflected, and the reflected coordinate was mapped back to the nearest voxel by rounding. Both source and reflected voxels had to lie within the analysis mask. Pearson correlations were clipped to ±0.999999 and Fisher transformed. This was a nearest-voxel mirror operation in MNI152NLin2009cAsym space; no additional symmetric-template registration was performed (Zuo et al., 2010).

#### Spatial smoothing and within-participant centring

For voxelwise analyses, the Nilearn 2-mm MNI grey-matter probability template was continuously resampled to the metric grid and thresholded at 0.20. Valid values were finite and positive for BEN, ALFF, fALFF and ReHo; signed VMHC maps required finite nonzero values. Gaussian smoothing with 8-mm full width at half maximum was applied separately to the masked map and its valid grey-matter support. Their ratio was retained where the smoothed support exceeded 0.05. The mean over each participant’s valid grey-matter support was then subtracted. We did not divide by the spatial standard deviation. The group mask required valid original-map coverage in at least 90% of participants and membership in the grey-matter mask.

Atlas-defined ROI analyses instead used complete valid cortical atlas support without an additional grey-matter threshold. Maps underwent analogous mask-normalized 8-mm smoothing and subtraction of the mean over valid cortical atlas support. ROI values were voxel-weighted means within the designated region. Thus, the voxelwise and ROI analyses had explicitly different spatial reference means.

#### Voxelwise statistical analysis

Ordinary least-squares models contained an intercept, duration CV, adjacent semantic similarity, age, sex and mean FD. Continuous predictors were standardized using sample standard deviations; the female indicator was mean-centred. Metric maps were spatially centred as described above, without subsequent voxelwise outcome standardization. The target coefficients were evaluated with two-sided t tests using residual degrees of freedom N − rank(X), equal to 69 in the base models. The central BEN contrast concerned duration CV; the complementary amplitude and synchronization contrasts concerned semantic similarity. Both behavioural predictors were included in each model.

Benjamini–Hochberg correction controlled FDR at q < 0.05 across tested voxels separately for each metric–behaviour coefficient map (Benjamini & Hochberg, 1995). It did not jointly control all coefficients in a regression or pool metrics and sensitivity models. No additional Bonferroni or across-metric correction was applied. Cluster sizes describe connected suprathreshold voxels, rather than cluster-level corrected tests. Model hierarchy describes the presentation of the completed analyses and does not imply preregistration.

#### Atlas-defined regional statistics

Cortical regions followed Schaefer’s 400-parcel atlas with 17-network assignments (Schaefer et al., 2018; Yeo et al., 2011). Default A, Default C and Control B were selected for correspondence with the original report; Default C here comprised cortical atlas parcels. Inferior parietal, retrosplenial and parahippocampal subdivisions followed label names. All valid voxels in each ROI were used, without restriction to significant voxels.

Models included semantic similarity, the speech-timing composite, articulation rate, transcript-event count, duration CV, age, sex and mean FD, plus an intercept (residual df = 66). Continuous predictors and regional outcomes were standardized; sex was mean-centred. We report nominal two-sided P values without correction across metrics or ROIs. Partial R² was t²/(t² + df). For the reported regional associations, sensitivity checks comprised HC3 inference, 10,000 Freedman–Lane permutations and leave-one-participant-out coefficient ranges. These regional tests were secondary, with subdivision localization exploratory.

A complementary parcelwise localization model used the same 75 participants and the same BEN processing, with sentence-event duration CV, adjacent semantic similarity, age, sex and mean FD as predictors. Parcel-mean BEN was extracted from all available Schaefer-400 parcels. Two-sided tests of the duration-CV coefficient were corrected across the 400 parcels using Benjamini–Hochberg FDR at q < 0.05. The full 400-parcel result is provided as Supplementary Data 1; the significant and near-threshold results are summarized in Supplementary Table 13.

#### Connectivity analysis

The contextual reanalysis extracted Schaefer parcel series from the fMRIPrep derivatives. An intercept, linear and quadratic trends, six motion parameters, global signal, white-matter signal and cerebrospinal-fluid signal were regressed from the full series. Missing confound entries were filled with zero. The first ten volumes and volumes with FD ≥ 1 mm together with two neighbours on either side were then excluded. No extra bandpass filter was used. Retained parcel series were standardized and averaged across parcels sharing a named bilateral subregion. Between-network connectivity was the mean Fisher-transformed correlation across subregion pairs; within-network connectivity used the upper triangle excluding the diagonal. Associations with thought-network clustering included age, sex and mean FD. This was a volumetric cortical approximation of the original analysis, including no hippocampal addition to Default C.

The new connectivity analysis used the BEN-matched 0.009–0.1 Hz series. The 20 FDR-significant BEN voxels were intersected with Default A and Control B atlas assignments to form 13- and 7-voxel seeds. Atlas labels and seeds were resampled with nearest-neighbour interpolation where required. Each seed series was the voxel mean. The target was the voxel mean across its corresponding network after excluding that seed, rather than an equal-weight average of parcel means. Pearson correlations were clipped to ±0.999999 and Fisher transformed. BEN values for the same seeds were extracted after grey-matter-confined 8-mm normalized smoothing and grey-matter centring.

#### Joint models and resampling

The outcome was standardized duration CV. The base design included standardized semantic similarity, age and mean FD and a centred female indicator. Nested models added standardized Control B seed BEN, seed-excluded Control B connectivity or both. Incremental R² was assessed with nested-model F tests. Percentile intervals used 20,000 participant bootstrap resamples with replacement (NumPy random seed 6067). The standardized design was resampled without repeating feature selection. Freedman–Lane tests used 10,000 permutations (seed 16067): reduced-model residuals were permuted, added to reduced-model fitted values, and the nested F statistic recomputed. P = (exceedances + 1)/(10,000 + 1). Joint coefficients also used HC3 standard errors and t-reference inference.

The seed was selected using duration CV in the same sample; bootstrap and permutation analyses held that seed fixed. Joint-model estimates therefore describe within-sample conditional associations. The fixed-feature leave-one-out prediction calculations in intermediate outputs are not used as independent validation in this article. Pearson correlation and regression, rather than rank correlations, underlie the reported analyses.

Repeated cross-validation used 20 repeats of five-fold partitioning of the same 75 participants (60 training, 15 test per fold). The base model contained adjacent semantic similarity, age, sex and mean FD. For each training fold, BEN voxelwise models were fit with those covariates; candidate voxels were selected either by the training-fold BH-FDR map or by a fixed, two-sided uncorrected p < 0.001 threshold, intersected with the Control B mask, and averaged to form one BEN feature. The fixed threshold was added after the initial FDR-based cross-validation and is therefore an exploratory sensitivity analysis. BEN and seed-excluded FC were compared with the covariate model using held-out Q² relative to the training-fold mean. Scaling and model fitting used training data only. Repeated splits reuse the same participants; ranges across repeats are split-sensitivity ranges, not confidence intervals or independent replications. We also summarized per-participant held-out error contributions and voxel-selection overlap across folds.

#### Temporal-unit sensitivity analysis

For each participant, we grouped sentence annotations by the released thoughtID and defined grouped-thought duration as the last sentence offset minus the first sentence onset, including intervening pauses. CV was the sample standard deviation divided by mean duration. Embedding counts and chronological order were checked against these groups. This CV replaced sentence-event CV in the same 75-participant models. To separate CV definition from count definition, count-adjusted models first retained sentence-event count; a further specification replaced it with the number of grouped thoughts. ROI contrasts followed the same sequence. Spatial processing and voxelwise correction were unchanged. With no FDR-significant grouped-thought BEN seed, the previous sentence-selected seeds were retained only for exploratory transfer analyses; no new seeds were defined from uncorrected maps. These analyses characterize sensitivity to operationalization, rather than a formal test of differences between temporal-level coefficients.

Annotation-level verification: all 75 BIDS event files matched the sentence annotation counts and onset/duration values within 0.011 s. Recomputed event-duration CV and adjacent embedding similarity matched the saved model values. Embedding row counts matched grouped thoughtID counts in every participant. The variable thought_count is retained in source tables for traceability and means sentence-level event count. Grouped-thought-duration CV is examined in the temporal-unit sensitivity analysis (Supplementary Tables 9–12). The new reanalysis table uses duration_cv and thought_count for grouped thoughts, and preserves sentence_duration_cv and sentence_event_count separately; the primary-analysis source files retain their historical sentence-event definitions.

### Supplementary Table 1 | Behavioural units and verification

N = 75. Counts and timing matched sentence annotations for all participants; embeddings matched grouped thoughts for all participants. Recomputed CV and semantic similarity matched saved analysis values. Individual checks are in Supplementary_behavioral_unit_audit.tsv.

| **Measure** | **Mean (SD)** | **Range** |
| --- | --- | --- |
| Sentence events / participant | 102.227 (35.659) | 41.000–206.000 |
| Grouped thoughts / participant | 54.747 (19.265) | 24.000–118.000 |
| Transcript-event duration CV | 0.711 (0.096) | 0.474–1.050 |
| Adjacent semantic similarity | 0.275 (0.050) | 0.104–0.387 |

### Supplementary Table 2 | Primary voxelwise FDR clusters

N = 75; two-sided OLS, base df = 69. BEN target: duration CV. Other metrics: adjacent semantic similarity. Each model includes the other behaviour, age, sex and FD. BH-FDR q < 0.05 was applied separately across voxels in each coefficient map. Cluster extent is descriptive. Maps use grey-matter-confined 8-mm smoothing and centring.

| **Metric** | **Sign** | **Voxels** | **Volume mm³** | **Peak t** | **MNI x, y, z** |
| --- | --- | --- | --- | --- | --- |
| BEN | negative | 13 | 96.7 | -5.632 | 27.5, 24.2, 54.0 |
| BEN | negative | 7 | 52.1 | -5.289 | 21.7, 41.6, 42.0 |
| ALFF | negative | 721 | 5365.7 | -7.351 | 60.2, -6.6, 24.0 |
| ALFF | negative | 546 | 4063.4 | -6.343 | -59.3, -6.6, 24.0 |
| ALFF | positive | 287 | 2135.9 | 5.125 | -22.7, -4.7, 68.0 |
| ALFF | positive | 80 | 595.4 | 5.319 | -36.2, 5.0, 54.0 |
| ALFF | negative | 57 | 424.2 | -4.312 | 35.2, -56.8, -20.0 |
| ALFF | positive | 47 | 349.8 | 4.284 | -30.4, 6.9, 54.0 |
| ALFF | negative | 29 | 215.8 | -4.081 | -65.1, -51.0, 10.0 |
| ALFF | positive | 27 | 200.9 | 4.152 | 14.0, 6.9, 42.0 |
| ALFF | positive | 20 | 148.8 | 3.864 | -45.8, -56.8, 30.0 |
| ALFF | negative | 15 | 111.6 | -4.108 | 64.1, -54.8, -12.0 |
| ALFF | positive | 12 | 89.3 | 3.782 | -24.6, -37.5, 6.0 |
| ALFF | positive | 10 | 74.4 | 4.106 | -36.2, 16.5, 46.0 |
| ALFF | positive | 7 | 52.1 | 3.963 | -18.8, -27.8, 56.0 |
| ALFF | positive | 7 | 52.1 | 3.898 | 6.2, -2.8, 56.0 |
| ALFF | positive | 6 | 44.7 | 3.813 | 10.1, 30.0, 38.0 |
| ALFF | positive | 6 | 44.7 | 3.809 | 27.5, -35.5, 54.0 |
| ALFF | positive | 5 | 37.2 | 3.711 | -32.3, 1.1, 14.0 |
| ALFF | negative | 4 | 29.8 | -3.752 | 15.9, -74.1, 36.0 |
| ALFF | positive | 4 | 29.8 | 3.729 | 21.7, -2.8, 66.0 |
| ALFF | positive | 3 | 22.3 | 3.693 | -13.1, 8.8, 12.0 |
| ALFF | positive | 2 | 14.9 | 3.744 | -15.0, -27.8, 48.0 |
| ALFF | positive | 2 | 14.9 | 3.675 | 14.0, 32.0, 26.0 |
| ALFF | positive | 2 | 14.9 | 3.651 | -3.4, -24.0, 58.0 |
| ALFF | positive | 2 | 14.9 | 3.716 | -26.6, 37.8, 30.0 |
| ALFF | positive | 1 | 7.4 | 3.638 | -20.8, -43.3, 58.0 |
| ALFF | positive | 1 | 7.4 | 3.681 | -18.8, -37.5, 68.0 |
| ALFF | positive | 1 | 7.4 | 3.738 | -11.1, -0.8, 52.0 |
| ALFF | positive | 1 | 7.4 | 3.632 | -9.2, 14.6, 58.0 |
| ALFF | negative | 1 | 7.4 | -3.630 | 14.0, -81.8, 46.0 |
| fALFF | negative | 657 | 4889.4 | -7.403 | 58.3, -6.6, 24.0 |
| fALFF | negative | 640 | 4762.9 | -7.078 | -53.6, -10.5, 28.0 |
| fALFF | positive | 157 | 1168.4 | 4.965 | -22.7, -2.8, 68.0 |
| fALFF | positive | 70 | 520.9 | 5.019 | -36.2, 5.0, 54.0 |
| fALFF | positive | 25 | 186.1 | 3.988 | -26.6, -16.3, 60.0 |
| fALFF | positive | 23 | 171.2 | 4.043 | -30.4, 6.9, 54.0 |
| fALFF | positive | 22 | 163.7 | 4.162 | 14.0, 6.9, 46.0 |
| fALFF | negative | 15 | 111.6 | -3.875 | -65.1, -51.0, 10.0 |
| fALFF | positive | 12 | 89.3 | 3.934 | -45.8, -56.8, 30.0 |
| fALFF | negative | 8 | 59.5 | -3.864 | 35.2, -56.8, -20.0 |
| fALFF | negative | 8 | 59.5 | -3.797 | 31.3, -85.7, 32.0 |
| fALFF | positive | 6 | 44.7 | 3.919 | -18.8, -27.8, 56.0 |
| fALFF | positive | 6 | 44.7 | 3.814 | -36.2, 16.5, 46.0 |
| fALFF | positive | 6 | 44.7 | 3.901 | 27.5, -35.5, 54.0 |
| fALFF | negative | 3 | 22.3 | -3.748 | 15.9, -74.1, 36.0 |
| fALFF | negative | 3 | 22.3 | -3.910 | 64.1, -54.8, -12.0 |
| fALFF | positive | 2 | 14.9 | 3.766 | 10.1, 30.0, 38.0 |
| fALFF | positive | 2 | 14.9 | 3.714 | 6.2, -4.7, 56.0 |
| fALFF | positive | 1 | 7.4 | 3.698 | -26.6, 37.8, 30.0 |
| fALFF | positive | 1 | 7.4 | 3.703 | -18.8, -37.5, 68.0 |
| fALFF | positive | 1 | 7.4 | 3.661 | -16.9, -24.0, 66.0 |
| fALFF | negative | 1 | 7.4 | -3.658 | -57.4, -49.1, 12.0 |
| fALFF | negative | 1 | 7.4 | -3.655 | -65.1, -16.3, 12.0 |
| ReHo | negative | 584 | 4346.2 | -6.164 | -55.5, -8.5, 26.0 |
| ReHo | negative | 499 | 3713.6 | -5.359 | 58.3, -0.8, 36.0 |
| ReHo | positive | 5 | 37.2 | 4.186 | -40.1, -72.2, 30.0 |
| ReHo | positive | 4 | 29.8 | 4.057 | 17.8, -35.5, -16.0 |
| VMHC | negative | 462 | 3438.2 | -6.124 | 62.2, -2.8, 20.0 |
| VMHC | negative | 435 | 3237.3 | -6.139 | -61.3, -2.8, 20.0 |
| VMHC | negative | 16 | 119.1 | -4.534 | 23.6, -62.6, -20.0 |
| VMHC | negative | 16 | 119.1 | -4.532 | -22.7, -62.6, -20.0 |
| VMHC | negative | 5 | 37.2 | -3.943 | -42.0, -52.9, -8.0 |
| VMHC | negative | 3 | 22.3 | -3.938 | 27.5, -10.5, 4.0 |

### Supplementary Table 3 | Speech-timing sensitivity models

FDR-significant voxel counts, N = 75 throughout. Separate BH-FDR q < 0.05 correction for each coefficient map. All models include duration CV, semantic similarity, age, sex and FD. Combined models add the indicated speech variables; event count is sentence-event count.

| **Model** | **ALFF** | **BEN** | **ReHo** | **VMHC** | **fALFF** |
| --- | --- | --- | --- | --- | --- |
| + Coverage | 0 | 40 | 0 | 0 | 0 |
| + Articulation rate | 1801 | 56 | 1175 | 961 | 1632 |
| + Mean pause | 0 | 40 | 0 | 0 | 0 |
| + Event count | 1945 | 20 | 1184 | 969 | 1753 |
| Mutual model | 1906 | 20 | 1092 | 937 | 1670 |
| Coverage + rate + count | 0 | 48 | 0 | 0 | 0 |
| Pause + rate + count | 0 | 31 | 0 | 0 | 0 |

### Supplementary Table 4 | Cortical ROI definitions

Schaefer 400 parcels / 17 networks. Complete valid atlas support, without an extra grey-matter threshold. Voxel counts refer to the ROI extraction grid. Full parcel labels are supplied in the source-data TSV. DefaultC_MTL_cortical is the full cortical Default C territory, not only its parahippocampal subdivision.

| **ROI** | **Parcels** | **Voxels** |
| --- | --- | --- |
| DefaultA_core | 34 | 12636 |
| DefaultC_MTL_cortical | 13 | 4080 |
| DefaultC_Rsp | 5 | 1231 |
| DefaultC_PHC | 5 | 1457 |
| DefaultC_IPL | 3 | 1392 |
| ControlB | 25 | 11406 |
| PMC_DefaultA_pCunPCC | 12 | 3276 |
| DefaultA_medial_PFC | 12 | 5241 |

### Supplementary Table 5 | Full cortical semantic models

N = 75; df = 66. Nominal two-sided tests, without across-metric or across-ROI correction. Predictors: semantic similarity, speech-timing contrast, articulation rate, event count, duration CV, age, sex and FD. Continuous variables and outcomes standardized. Intervals are conventional OLS coefficient intervals. Network results are secondary; subdivisions and additional cortical regions exploratory.

| **ROI** | **Metric** | **β** | **95% CI** | **P** |
| --- | --- | --- | --- | --- |
| DefaultA_core | BEN | 0.111 | -0.221, 0.443 | 0.506 |
| DefaultC_MTL_cortical | BEN | 0.162 | -0.201, 0.524 | 0.376 |
| DefaultC_Rsp | BEN | 0.189 | -0.175, 0.553 | 0.303 |
| DefaultC_PHC | BEN | 0.049 | -0.325, 0.424 | 0.793 |
| DefaultC_IPL | BEN | 0.091 | -0.277, 0.460 | 0.623 |
| ControlB | BEN | -0.159 | -0.513, 0.195 | 0.372 |
| PMC_DefaultA_pCunPCC | BEN | 0.203 | -0.158, 0.564 | 0.266 |
| DefaultA_medial_PFC | BEN | 0.017 | -0.326, 0.359 | 0.922 |
| DefaultA_core | ALFF | 0.100 | -0.259, 0.459 | 0.579 |
| DefaultC_MTL_cortical | ALFF | -0.021 | -0.383, 0.342 | 0.909 |
| DefaultC_Rsp | ALFF | -0.201 | -0.558, 0.156 | 0.264 |
| DefaultC_PHC | ALFF | -0.028 | -0.393, 0.337 | 0.878 |
| DefaultC_IPL | ALFF | 0.156 | -0.209, 0.521 | 0.396 |
| ControlB | ALFF | 0.081 | -0.271, 0.434 | 0.647 |
| PMC_DefaultA_pCunPCC | ALFF | -0.028 | -0.387, 0.330 | 0.875 |
| DefaultA_medial_PFC | ALFF | 0.097 | -0.265, 0.458 | 0.595 |
| DefaultA_core | fALFF | 0.102 | -0.257, 0.460 | 0.574 |
| DefaultC_MTL_cortical | fALFF | -0.049 | -0.409, 0.311 | 0.786 |
| DefaultC_Rsp | fALFF | -0.227 | -0.578, 0.123 | 0.2 |
| DefaultC_PHC | fALFF | -0.047 | -0.410, 0.316 | 0.795 |
| DefaultC_IPL | fALFF | 0.140 | -0.229, 0.509 | 0.451 |
| ControlB | fALFF | 0.088 | -0.263, 0.438 | 0.619 |
| PMC_DefaultA_pCunPCC | fALFF | -0.079 | -0.432, 0.275 | 0.659 |
| DefaultA_medial_PFC | fALFF | 0.121 | -0.242, 0.484 | 0.509 |
| DefaultA_core | ReHo | -0.121 | -0.487, 0.244 | 0.51 |
| DefaultC_MTL_cortical | ReHo | 0.517 | 0.180, 0.854 | 0.00316 |
| DefaultC_Rsp | ReHo | 0.103 | -0.256, 0.463 | 0.568 |
| DefaultC_PHC | ReHo | 0.282 | -0.075, 0.639 | 0.12 |
| DefaultC_IPL | ReHo | 0.499 | 0.153, 0.846 | 0.00536 |
| ControlB | ReHo | 0.028 | -0.315, 0.372 | 0.87 |
| PMC_DefaultA_pCunPCC | ReHo | -0.089 | -0.406, 0.228 | 0.577 |
| DefaultA_medial_PFC | ReHo | -0.140 | -0.506, 0.226 | 0.449 |
| DefaultA_core | VMHC | -0.042 | -0.395, 0.311 | 0.814 |
| DefaultC_MTL_cortical | VMHC | 0.281 | -0.062, 0.624 | 0.106 |
| DefaultC_Rsp | VMHC | -0.213 | -0.592, 0.166 | 0.265 |
| DefaultC_PHC | VMHC | 0.316 | -0.035, 0.667 | 0.0769 |
| DefaultC_IPL | VMHC | 0.412 | 0.082, 0.741 | 0.0151 |
| ControlB | VMHC | -0.020 | -0.400, 0.360 | 0.917 |
| PMC_DefaultA_pCunPCC | VMHC | -0.056 | -0.378, 0.266 | 0.73 |
| DefaultA_medial_PFC | VMHC | 0.003 | -0.372, 0.378 | 0.986 |

### Supplementary Table 6 | Regional robustness checks

Same N = 75 model as Supplementary Table 5. HC3 heteroscedasticity-robust tests; 10,000 Freedman–Lane permutations; leave-one-participant-out (LOO) coefficient ranges. All tests nominal. The table includes the additional Default C parahippocampal VMHC check in the saved robustness output.

| **ROI / metric** | **β** | **P OLS** | **P HC3** | **P perm.** | **LOO β range** |
| --- | --- | --- | --- | --- | --- |
| DefaultC_MTL_cortical / ReHo | 0.517 | 0.00316 | 0.00998 | 0.003 | 0.398, 0.600 |
| DefaultC_IPL / ReHo | 0.499 | 0.00536 | 0.0216 | 0.0059 | 0.406, 0.633 |
| DefaultC_IPL / VMHC | 0.412 | 0.0151 | 0.0129 | 0.0163 | 0.336, 0.481 |
| DefaultC_PHC / VMHC | 0.316 | 0.0769 | 0.167 | 0.0765 | 0.250, 0.465 |

### Supplementary Table 7 | Contextual connectivity reanalysis

N = 75; residual df = 70. Atlas-based cortical approximation of the original pipeline. Associations use thought-network clustering and age, sex and FD. This semantic graph measure differs from adjacent semantic similarity. Nominal tests; full source schema is preserved in Supplementary_Table7_original_FC_full.tsv.

| **outcome** | **beta** | **t** | **p** |
| --- | --- | --- | --- |
| Control B–Default A | 0.261 | 2.316 | 0.0235 |
| Within cortical Default C | -0.252 | -2.158 | 0.0343 |

### Supplementary Table 8 | Joint-model incremental variance

N = 75. Outcome: standardized transcript-event duration CV. Base covariates: semantic similarity, age, sex and FD. BEN and FC refer to the Control B prefrontal seed and its seed-excluded target. Bootstrap: 20,000 resamples, seed 6067. Freedman–Lane: 10,000 permutations, seed 16067. Feature selection was fixed; estimates are within sample. Nonnegative ΔR² bootstrap intervals are not used to determine significance.

| **Addition** | **ΔR²** | **F** | **df** | **P** | **Bootstrap 95% interval** | **P perm.** |
| --- | --- | --- | --- | --- | --- | --- |
| BEN beyond base | 0.274 | 31.207 | 1, 69 | 4.28e-07 | 0.1174, 0.4325 | 0.0001 |
| FC beyond base | 0.076 | 6.560 | 1, 69 | 0.0126 | 0.0049, 0.1980 | 0.0134 |
| BEN beyond FC | 0.231 | 27.450 | 1, 68 | 1.71e-06 | 0.0813, 0.3998 | 0.0001 |
| FC beyond BEN | 0.033 | 3.973 | 1, 68 | 0.0502 | 0.0004, 0.1171 | 0.0524 |
| BEN + FC beyond base | 0.307 | 18.262 | 2, 68 | 4.49e-07 | 0.1652, 0.4536 | 0.0001 |

### Supplementary Table 9 | Temporal-unit replacement in the mutual model

N = 75; df = 69. FDR-significant voxel counts. BEN target: duration CV; all other targets: semantic similarity. Models include the other behavioural variable, age, sex and FD. No count covariate is included in these mutual base models. Separate voxelwise BH-FDR q < 0.05 per coefficient map. All seven grouped-thought specifications in the main reanalysis and the three fixed-sentence-count specifications had zero BEN FDR voxels.

| **Metric** | **Sentence CV** | **Thought CV** |
| --- | --- | --- |
| BEN | 20 | 0 |
| ALFF | 1906 | 1600 |
| fALFF | 1670 | 1423 |
| ReHo | 1092 | 923 |
| VMHC | 937 | 563 |

### Supplementary Table 10 | Separating duration and count definitions

N = 75. FDR-significant voxel counts. Read left to right: replace CV while holding sentence count fixed, then replace count while holding thought CV fixed. Underlying imaging, behavioural mutual adjustment, age, sex, FD and per-map BH-FDR remain identical. Count-only df = 68; combined df = 66. Complete maps and model summaries are archived separately.

| **Metric** | **Added terms** | **Sentence CV / sentence count** | **Thought CV / sentence count** | **Thought CV / thought count** |
| --- | --- | --- | --- | --- |
| BEN | + Count | 20 | 0 | 0 |
| BEN | + Coverage, rate, count | 48 | 0 | 0 |
| BEN | + Pause, rate, count | 31 | 0 | 0 |
| ALFF | + Count | 1945 | 1673 | 1548 |
| ALFF | + Coverage, rate, count | 0 | 0 | 0 |
| ALFF | + Pause, rate, count | 0 | 0 | 0 |
| fALFF | + Count | 1753 | 1521 | 1454 |
| fALFF | + Coverage, rate, count | 0 | 0 | 0 |
| fALFF | + Pause, rate, count | 0 | 0 | 0 |
| ReHo | + Count | 1184 | 994 | 957 |
| ReHo | + Coverage, rate, count | 0 | 0 | 0 |
| ReHo | + Pause, rate, count | 0 | 0 | 0 |
| VMHC | + Count | 969 | 605 | 581 |
| VMHC | + Coverage, rate, count | 0 | 0 | 0 |
| VMHC | + Pause, rate, count | 0 | 0 | 0 |

### Supplementary Table 11 | Semantic ROI associations across temporal definitions

N = 75; df = 66. Definition column gives CV level / count level. Models also include speech-timing contrast, articulation rate, age, sex and FD. Standardized coefficients; conventional 95% intervals; nominal two-sided P values without across-ROI or across-metric correction. Full cortical tables and HC3 results are included in source data.

| **ROI / metric** | **CV / count** | **β** | **95% CI** | **P** |
| --- | --- | --- | --- | --- |
| Default C ReHo | Sentence / sentence | 0.517 | 0.180, 0.854 | 0.003155 |
| Default C ReHo | Thought / sentence | 0.501 | 0.184, 0.817 | 0.002431 |
| Default C ReHo | Thought / thought | 0.511 | 0.204, 0.818 | 0.001453 |
| IPL ReHo | Sentence / sentence | 0.499 | 0.153, 0.846 | 0.005361 |
| IPL ReHo | Thought / sentence | 0.469 | 0.143, 0.794 | 0.005418 |
| IPL ReHo | Thought / thought | 0.459 | 0.142, 0.777 | 0.00527 |
| IPL VMHC | Sentence / sentence | 0.412 | 0.082, 0.741 | 0.01512 |
| IPL VMHC | Thought / sentence | 0.500 | 0.184, 0.816 | 0.002371 |
| IPL VMHC | Thought / thought | 0.474 | 0.166, 0.782 | 0.003094 |

### Supplementary Table 12 | Exploratory transfer of the previous Control B seed

N = 75. Outcome: grouped-thought duration CV. Base covariates: semantic similarity, age, sex and FD. Seed BEN and strictly seed-excluded FC are identical to the sentence-level analysis. The previous sentence-selected seed is held fixed. Nominal nested F tests and 10,000 Freedman–Lane permutations; full 20,000-resample bootstrap intervals are in source data. These are exploratory transfer estimates, not a new grouped-thought seed discovery.

| **Added term** | **ΔR²** | **F** | **df** | **P** | **P perm.** |
| --- | --- | --- | --- | --- | --- |
| BEN beyond base | 0.0054 | 0.402 | 1, 69 | 0.5282 | 0.5371 |
| FC beyond base | 0.0007 | 0.049 | 1, 69 | 0.8256 | 0.8240 |
| BEN beyond FC | 0.0049 | 0.359 | 1, 68 | 0.5511 | 0.5477 |
| FC beyond BEN | 0.0002 | 0.011 | 1, 68 | 0.9161 | 0.9174 |
| BEN + FC beyond base | 0.0056 | 0.204 | 2, 68 | 0.8162 | 0.8197 |

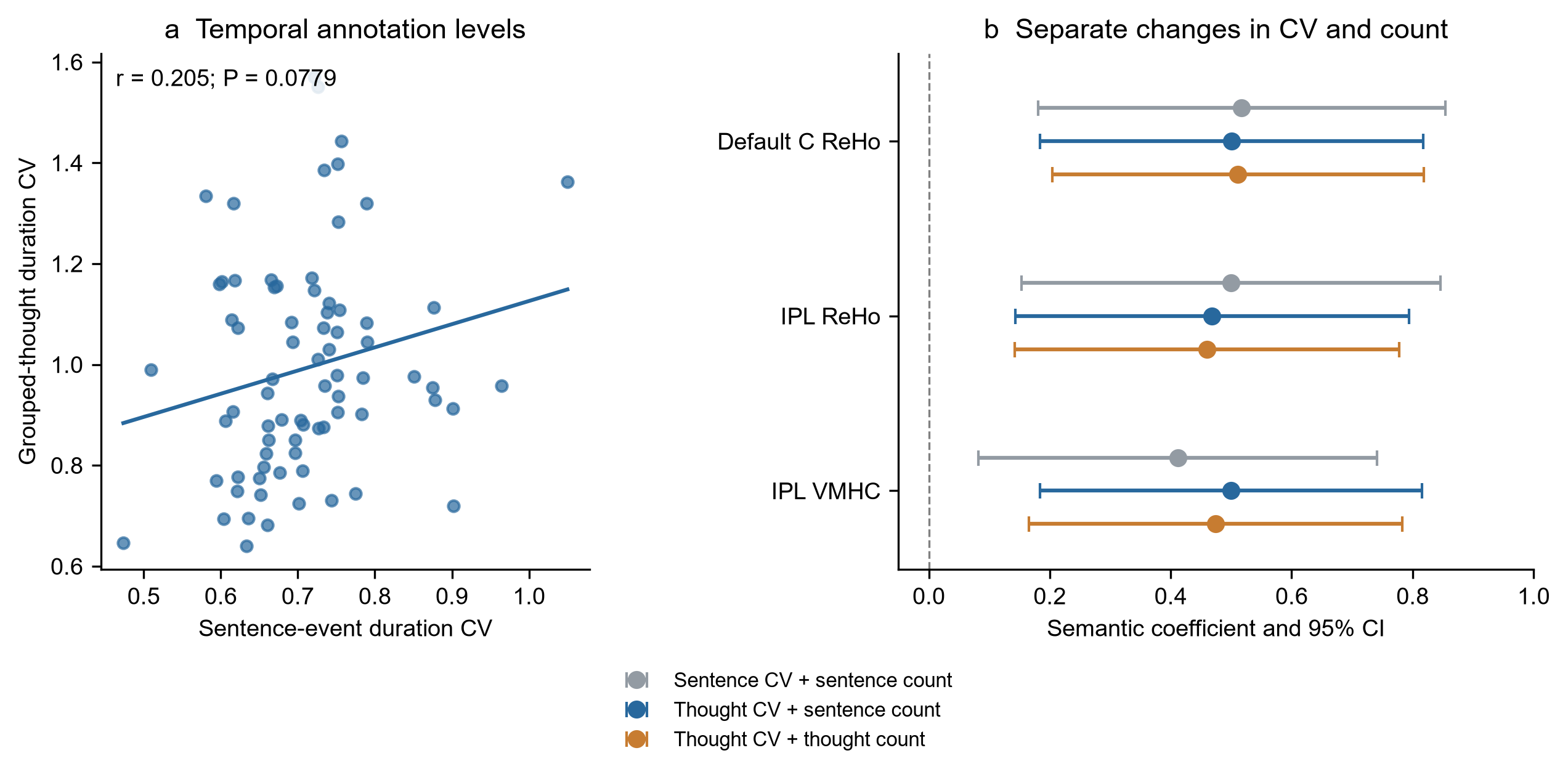

Supplementary Figure 1 | Temporal-unit and count-definition sensitivity. a, Sentence-event and grouped-thought duration CV in the same 75 participants. Each dot represents a participant; the line is an OLS fit. Pearson r = 0.205, two-sided P = 0.0779. b, Standardized semantic coefficients and conventional 95% confidence intervals from the three speech-adjusted ROI associations. Grey: sentence CV and sentence count. Blue: thought CV with sentence count held fixed. Orange: thought CV and thought count. All models include the speech-timing contrast, articulation rate, age, sex and FD (df = 66); tests are nominal. The panel describes sensitivity to variable definitions and does not test between-model differences. Source data are provided.

### Supplementary Table 13 | Parcelwise BEN associations with sentence-event duration variability

N = 75. Standardized parcel-mean BEN outcomes and standardized continuous predictors; each model includes adjacent semantic similarity, age, sex and mean FD. Two-sided p values for sentence-event duration CV were corrected across 400 Schaefer parcels using BH-FDR. All significant parcels and parcels with q < 0.10 are shown; the complete 400-parcel table is supplied as Supplementary Data 1.

| **Schaefer parcel** | **Network** | **Std. β** | **t(69)** | **P** | **q (400)** |
| --- | --- | --- | --- | --- | --- |
| RH_DefaultA_PFCd_1 | DefaultA | -0.453 | -4.009 | 0.000152 | 0.0310 |
| RH_DefaultB_PFCd_3 | DefaultB | -0.455 | -3.951 | 0.000185 | 0.0310 |
| RH_ContB_PFCld_1 | ContB | -0.450 | -3.884 | 0.000232 | 0.0310 |
| RH_ContB_PFCld_2 | ContB | -0.432 | -3.775 | 0.000335 | 0.0335 |
| RH_DefaultB_PFCd_4 | DefaultB | -0.401 | -3.543 | 0.000714 | 0.0534 |
| RH_ContA_PFCl_2 | ContA | -0.401 | -3.487 | 0.000853 | 0.0534 |
| RH_DefaultA_PFCd_2 | DefaultA | -0.395 | -3.459 | 0.000934 | 0.0534 |
| LH_SalVentAttnB_PFCl_3 | SalVentAttnB | -0.400 | -3.389 | 0.00116 | 0.0582 |

### Supplementary Table 14 | Repeated cross-validation performance under alternative feature-selection rules

Outcome: sentence-event duration CV. Baseline covariates: adjacent semantic similarity, age, sex and mean FD. Each value summarizes 20 repeated five-fold partitions of the same N = 75 cohort. Repeat range is split sensitivity, not a confidence interval. The fixed p threshold was introduced after inspecting the training-fold FDR analysis.

| **Feature selection** | **Nonempty folds** | **Mean ΔQ²** | **Repeat range** | **Positive repeats** | **Note** |
| --- | --- | --- | --- | --- | --- |
| Training-fold BH-FDR | 28/100 | −0.001 | −0.092 to 0.163 | 9/20 | Seed sometimes empty |
| Training-fold p < 0.001 | 100/100 | 0.080 | −0.017 to 0.188 | 19/20 | Post hoc fixed threshold; median 92 voxels |
| Training-fold p < 0.005 | 100/100 | 0.064 | −0.044 to 0.166 | 18/20 | Sensitivity; median 317 voxels |
| Training-fold p < 0.01 | 100/100 | 0.056 | −0.093 to 0.153 | 17/20 | Sensitivity; median 514 voxels |

### Supplementary Table 15 | Participant influence and spatial stability of fixed-threshold BEN features

Positive participant contributions indicate lower held-out squared error after adding BEN than in the corresponding covariate model. These are contributions to the existing held-out scores, not full leave-one-participant-out retraining. Jaccard values summarize pairwise overlap among the 100 training-fold voxel selections.

| **Training-fold threshold** | **Positive contributors** | **Largest single contribution** | **Median Jaccard** | **Voxels selected in ≥50% folds** |
| --- | --- | --- | --- | --- |
| p < 0.001 | 40/75 | 79.8% | 0.258 | 59 |
| p < 0.005 | 38/75 | 99.9% | 0.328 | 238 |
| p < 0.01 | 36/75 | >100% of net gain | 0.378 | 416 |

### Supplementary Figure 2 | ALFF associations and correspondence with fALFF

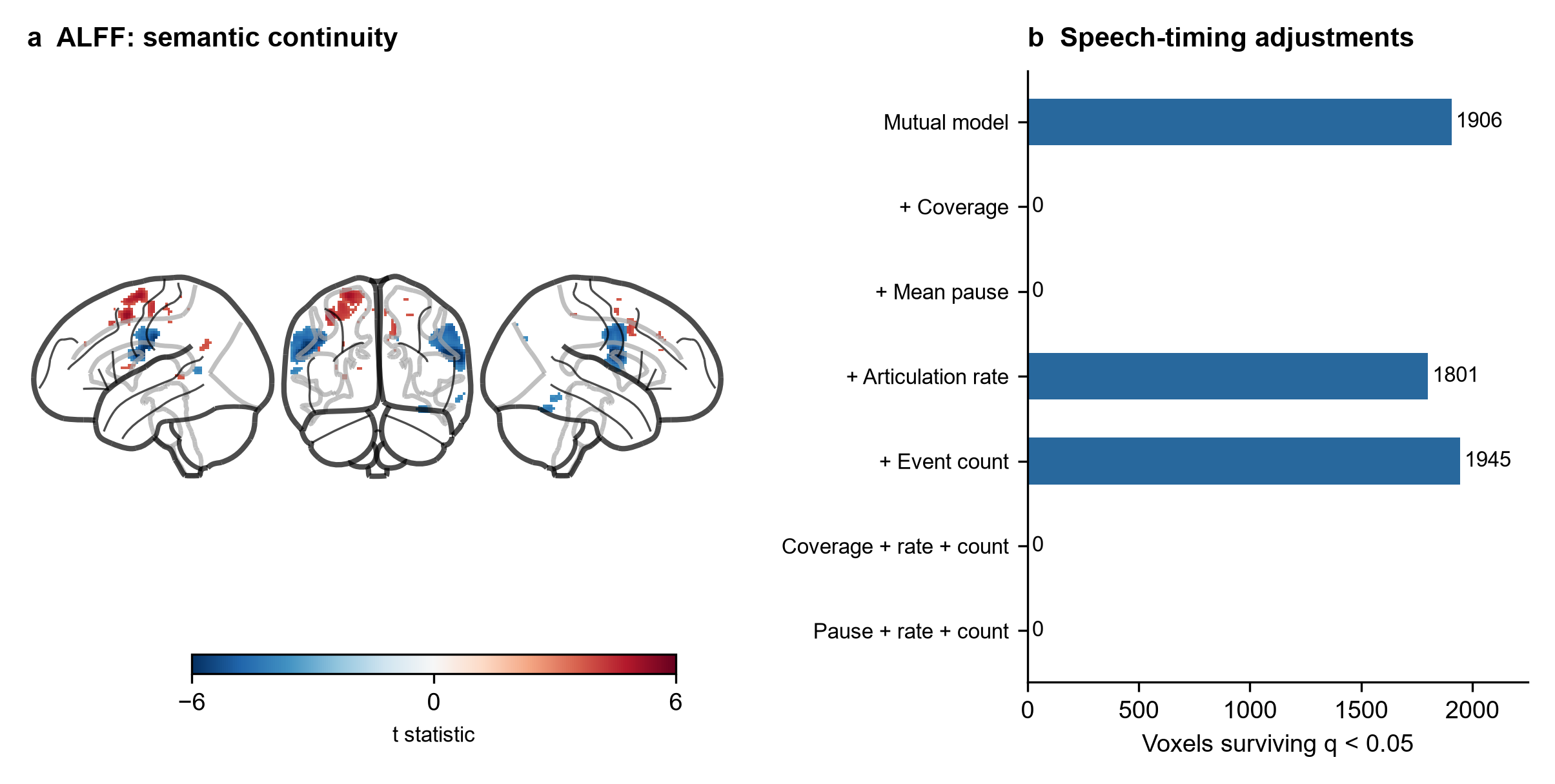

a, Signed glass-brain projections of the ALFF semantic-similarity coefficient after voxelwise BH-FDR q < 0.05. Models include sentence-event duration CV, age, sex and mean FD (N = 75; base df = 69). Blue and red indicate negative and positive coefficients. Maps use grey-matter-confined 8-mm smoothing and within-participant centring. b, Significant voxel counts after individual and combined speech-timing adjustments. Each coefficient map is corrected separately across voxels. Counts describe thresholded maps and are not tests of differences between models.

ALFF and fALFF shared 1,549 FDR-significant voxels: 81.3% of the 1,906 ALFF voxels and 92.8% of the 1,670 fALFF voxels (Dice = 0.866). Coefficient signs agreed in all shared voxels. Both analyses yielded no FDR-significant voxels after adjustment for speech coverage or mean pause. ALFF retained 1,801 voxels after articulation-rate adjustment and 1,945 after sentence-count adjustment; the corresponding fALFF counts were 1,632 and 1,753. With grouped-thought duration CV, ALFF retained 1,600 voxels in the mutual model. These comparisons describe spatial and model-adjustment concordance; they do not establish metric equivalence. Full ALFF ROI and sensitivity results are retained in the supplementary tables and source data.
